# Joint Estimation of Mother and Daughter Protein Regulatory Networks Quantifies Inheritance in Yeast Lineages

**DOI:** 10.64898/2026.11.20.767944

**Authors:** Shirley Mathur, Wenbin Wu, Taylor Kennedy, Orlando Argüello-Miranda, Kevin Z. Lin

## Abstract

Live-cell imaging tracks multiple protein levels within individual cells across divisions in real time, enabling direct measurement of how regulatory dynamics are inherited by a daughter cell from her mother. This technology presents a unique opportunity to directly observe longitudinal dynamics, in contrast to single-cell RNA sequencing, which can only measure cells cross-sectionally. However, there is a lack of computational methods for analyzing live-cell imaging data. Existing frameworks based on ordinary differential equations (ODEs) estimate networks from individual time series, but adapting them to mother-daughter pairs poses a dilemma: pooling both cells forces complete inheritance, while fitting each cell separately starves the short daughter trajectory of timepoints needed for reliable estimation. To address this gap, we develop PRNODE (Protein Regulatory Networks via ODEs), which models each cell’s network as a deviation from a shared regulatory scaffold and estimates the scaffold and both deviations jointly under a fused group-lasso penalty. The mother’s longer time series thereby supports the daughter’s estimate, while allowing the two networks to differ wherever the data demand it. We then propose a novel measure of inheritance based on comparing the rate of change of protein levels between the mother’s and daughter’s networks. Across simulation studies, we find that PRNODE outperforms existing ODE estimation methods in recovering the networks of daughter cells and in estimating inheritance. Finally, we apply PRNODE to many mother-daughter pairs of *Saccharomyces cerevisiae* observed over a 9-hour period at 12-minute intervals. Many estimated networks show a negative feedback loop between the cell-cycle inhibitor Whi5 and the cyclin Clb2, which together sustain cell-cycle oscillations. We also find that the measured inheritance tracks the number of regulatory edges a daughter shares with her mother with the same sign, more than the presence of any single edge. PRNODE is available as an R package at https://github.com/shirmath/LIC_methods.

## 1 Introduction

Advances in fluorescent reporters, microfluidics, and deep-learning-based segmentation and tracking now make it possible to follow an individual cell through several divisions and to record the abundance of multiple proteins in that cell at every timepoint [2,20,10,18]. The resulting live-cell imaging data are a multivariate time series for each cell together with the observed lineage that links each daughter to her mother (Figure 1A and B), so that both the time of each measurement and the relationship between cells are directly measured rather than inferred. In contrast, single-cell RNA sequencing, a common and dominant source of cell-level data, destroys each cell it measures, so that its analyses infer each cell’s position in time and relate cells only at the times they are sequenced (Table 1) [37,13,32,41,36]. Live-cell imaging has revealed that genetically identical cells in the same culture differ in their protein dynamics and in the states they enter [3], and it makes it possible to ask what a dividing cell passes on to its daughter. We develop **PRNODE** (Protein Regulatory Networks via Ordinary Differential Equations) to advance the computational methods for understanding protein regulatory dynamics in live-cell imaging data (Figure 1C). Our method estimates the protein regulatory networks of a mother cell and one of its daughters jointly, allowing the two networks to differ, and quantifies how much of the mother’s network the daughter inherits. Measuring inheritance at the level of regulatory mechanism, rather than of individual protein abundances, asks how a cell transmits its state to the next generation, the question underlying cellular memory of stress in yeast and lineage-dependent fates in mammalian cells [3,27].

**Table 1.** What each source of cell-level data observes, and the unit for which a regulatory network can be estimated from it. This paper: (*)

| Data (typical analysis) | Time of a measurement | Lineage | Measured | Network for |
|---|---|---|---|---|
| scRNA-seq (pseudotime, velocity) [37,13] | Inferred | None | Transcriptome | Population |
| scRNA-seq with static lineage barcodes [36,19] | Sampling times | Clones at sequencing | Transcriptome | Population at clone-level |
| Time-course scRNA-seq [41,17] | Per sample, not per cell | None | Transcriptome | Population |
| **Live-cell imaging**<sup>(\*)</sup> [20,18] | Observed, per cell | Mother-daughter | A few proteins | Individual cells |

**Figure 1.**
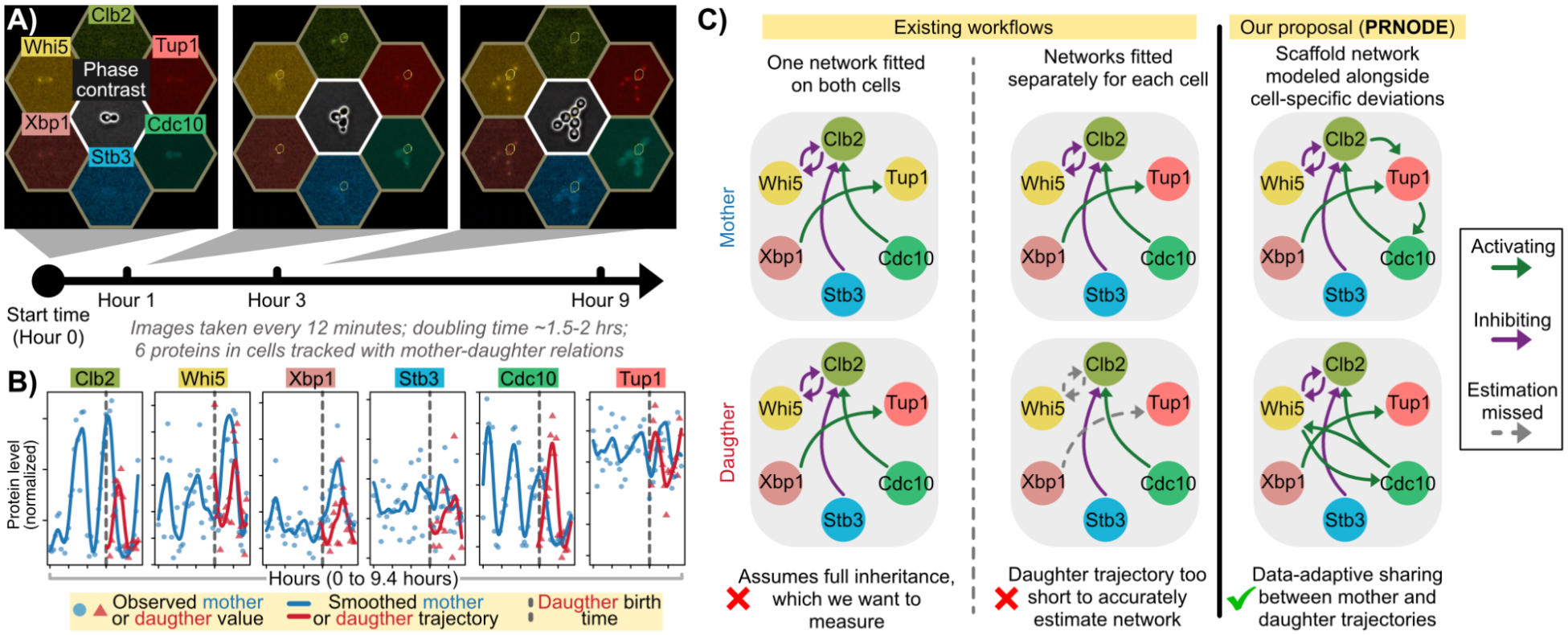
Live-cell imaging data and motivation of PRNODE. **A**) Schematic of live-cell imaging data where cells are longitudinally tracked across 6 proteins alongside the phase contrast. Each bordering hexagon illustrates the imaging detecting abundance of a particular fluorescent protein binding to a particular protein. **B**) Deep-learning methods allow us to segment, track, and extract numerical features from each protein in every cell. The results are time-series data such as the example shown here, where we visualize the measured values (dots) and smoothed trajectories. **C**) Premise of our paper, demonstrating how PRNODE overcomes gaps present in existing workflows.

Our analysis uses budding yeast (*Saccharomyces cerevisiae*), whose time between divisions is under two hours in rich medium, short enough that one experiment records several generations of a lineage. Live-cell imaging of yeast has already established that a daughter cell is shaped by its mother beyond their shared genome. Transcription factors and organelles are partitioned asymmetrically at division [14,15], daughter-specific transcription factors set a daughter’s size control apart from its mother’s [12], and correlated division times across lineages of bacterial and mammalian cells point to latent factors passed between generations [27,25,16]. Most directly, Argüello-Miranda et al. [2] track six proteins in individual starving yeast cells and show that whether a daughter will enter meiosis or quiescence is predictable from its mother’s biomarkers before the daughter is born. These results suggest that part of the mother’s regulatory state passes to the daughter, but which components of that machinery are transmitted at division, and which are reset, remains unknown.

The yeast cell cycle is driven by a network of proteins regulating one another, understood well enough that ODE models of it reproduce the phenotypes of over a hundred mutant strains [7,35]. A protein regulatory network records, for each protein, how the abundances of the other proteins raise or lower its rate of change over time, so that the network together with the current protein levels determines the trajectories that follow. Among the proteins we image, for example, nuclear Whi5 holds the cell in G1 and the mitotic cyclin Clb2 drives it through mitosis, a loop of opposite phases described in Section 5 and Appendix G [33,1,7]. Because the network determines how a cell’s protein levels evolve, a daughter that inherits its mother’s network inherits the dynamics that produced its mother’s behavior, even when asymmetric partitioning leaves the daughter with different initial protein levels than its mother [15]. We therefore study inheritance as the similarity between the mother’s regulatory network and the daughter’s, which requires estimating both networks from the imaging data. Sparse ODE methods such as GRADE estimate which proteins regulate which from noisy, discretely sampled trajectories [6,38,8,11], but they fit either one network to a population of time series or one network per series. Multi-task methods share networks across the cell types of a lineage tree [42,29], but their units are cell types observed in snapshots rather than two cells with observed time series and an observed division between them. Neither setting describes a mother-daughter pair, whose two cells should share much of their network but not all of it, and whose daughter, born partway through the experiment, has far fewer timepoints than her mother. PRNODE extends GRADE to this setting by estimating the two networks jointly under a fused penalty that shrinks both toward a shared network, so that the mother’s longer series supports the daughter’s estimate while the two networks are allowed to differ where the data demand it. How similarly the two estimated networks move a cell’s protein levels over one sampling interval is then our measure of inheritance.

## 2 Background

Live-cell imaging observes both the time course of each cell and its lineage, which pseudotime, RNA velocity, and lineage barcodes must infer from single-cell RNA sequencing [37,13,36] (Table 1, Appendix A), at the cost of measuring a few proteins rather than the transcriptome, so that a regulatory network can be estimated for each cell and compared between a mother and her daughter. Estimating such a network requires a model of how the abundance of each protein changes the others’ over time, which we set up for the two cells of a mother-daughter pair at once. A cell *c* ∈ *{M, D}*, the mother or her daughter, is imaged at *n* equally spaced timepoints *t*_1_ *< · · · < t*_*n*_ on *p* proteins, with time rescaled so that the mother’s series spans [0, 1] and the daughter, born at 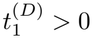, is observed at the same timepoints until the end of the experiment. The measured value of protein *j* at the *i*-th timepoint of cell *c* is modeled as 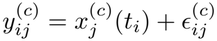, where 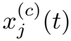 is the noiseless abundance and the errors 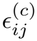 are independent with mean zero and variance 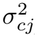. The standard model for the noiseless abundances is a system of ordinary differential equations (ODEs), 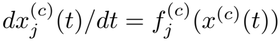, in which protein *k* regulates protein *j* when 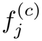 depends on 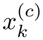 [4,8], so that estimating the network amounts to estimating which proteins each 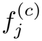 depends on. Sparse identification of nonlinear dynamics (SINDy) writes each *f*_*j*_ as a sparse combination of candidate functions and regresses a numerical derivative of *x*_*j*_ on them [5], and weak formulations avoid the noisy derivative by integrating against smooth test functions [26,24]. GRADE [8] instead assumes that each 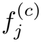 is additive and approximates each additive component in the polynomial basis *ψ*(*x*) = [*x, x*^2^, …, *x*^*q*^]^⊤^,

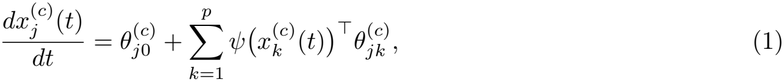

where 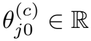 is the constant term, and 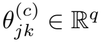 contains the coefficients of regulator *k* in the equation of protein *j* (Figure 2A). The network of cell *c* is the adjacency matrix 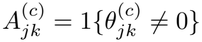, with an edge *k → j* when protein *k* enters the rate equation of protein *j*. Because (1) is linear in its coefficients, integrating it from the cell’s first timepoint allows us to model the trajectory itself, up to the integrated approximation error, as a linear function of the same coefficients,

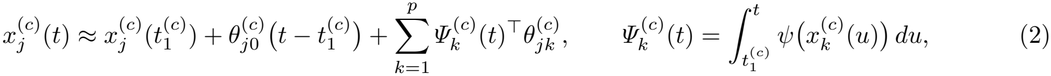

for any 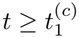. Therefore, the observed series can be regressed on the integrated basis 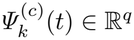 of each regulator without differentiating noisy data, while the initial condition and the constant term multiply the known functions 1 and 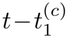. GRADE [8] computes 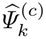 from a smoothing-spline estimate of each trajectory and fits (2) to one cell at a time by least squares with a group-lasso penalty [40] on the fitted contribution 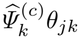 of each regulator [30], so that the *q* coefficients of regulator *k* leave the equation together and the estimated network has an edge *k → j* exactly when 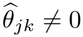. See Appendix A for more discussion on SINDy and GRADE.

**Figure 2.**
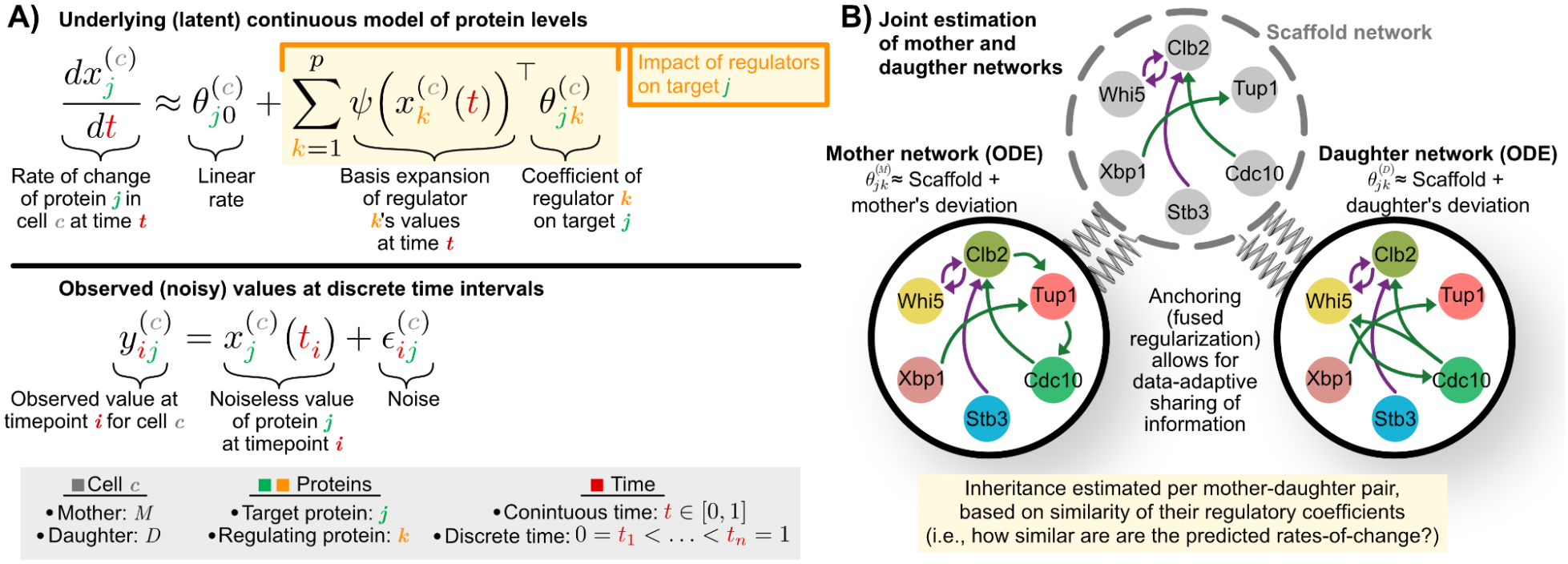
Notation and PRNODE schematic. **A**) Notation scheme to model the interplay between protein values within individual cell across time. **B**) Conceptual illustration of PRNODE, where the scaffold network represents the network structure shared between both the mother and daughter cells while cell-specific deviations are allowed.

The alternatives to GRADE fit these data poorly for three reasons: pseudotime and velocity methods applied to scRNA-seq data must infer the time and lineage in contrast to live-cell imaging’s direct measurements, Granger-style methods infer regulation from the order of a population of cells rather than from one cell’s trajectory [32,31], and a neural ODE [41,17] has far more parameters than the at most 48 timepoints of one cell, imaged every 12 minutes, can estimate. Kernel ODE (KODE) [11] replaces GRADE’s polynomial basis by a reproducing-kernel expansion of each additive component, and our earlier analysis of live-cell yeast data fits it to each cell on its own [39], so that the daughter’s network rests on her short series alone. GRADE fits one cell well only when the cell has many timepoints, yet a daughter, born partway through her mother’s series, has fewer timepoints than her mother, and pooling the two through the multiple-series form of GRADE forces them to share one network, which assumes the complete inheritance that the analysis sets out to measure. An estimator that borrows the mother’s timepoints where the two networks agree and lets them differ elsewhere should estimate the daughter’s network, and with it the inheritance, more accurately, which motivates PRNODE.

## 3 Methods

### 3.1 The PRNODE estimator

PRNODE estimates the mother’s and the daughter’s networks jointly, so that the mother’s additional data points can help improve estimation of the daughter’s network. We introduce a scaffold coefficient 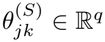 for each pair of proteins and write each cell’s coefficients in (1) as the scaffold plus a deviation,

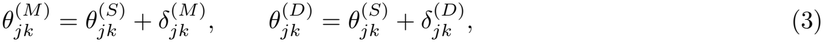

so that the scaffold reflects the regulatory structure shared by both cells and a deviation 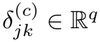 accounts for edges specific to cell *c* or changes in the strength of a shared edge (Figure 2B). We formulate the loss objective as

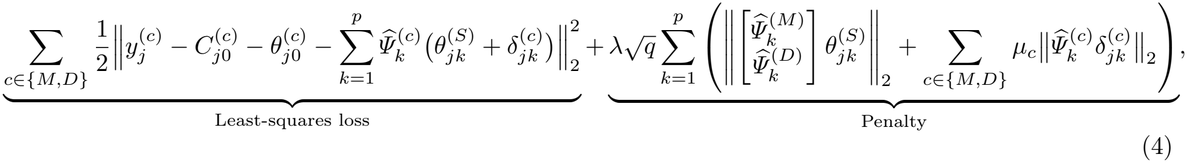

where 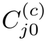 is the initial condition of protein *j* for cell *c*, 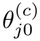 is the constant term in the linear ODE approximation in (2), and *λ, µ*_*M*_, *µ*_*D*_ are tuning parameters. The first term in the loss corresponds to the least-squares loss of the estimated model for the observed protein intensities, and the second term corresponds to penalizing the coefficients governing the structure of the regulatory networks (i.e., all ODE coefficients aside from the initial condition and the solely time-dependent term of the ODE). Since 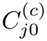 and 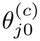 are not penalized in (4), estimating these parameters as the values minimizing the loss has a closed-form solution. Thus, we instead use a profiled form of the loss to estimate the penalized coefficients, where for each protein *j* we estimate 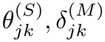, and 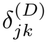 for *k* ∈ *{*1, …, *p}* as the minimizers of

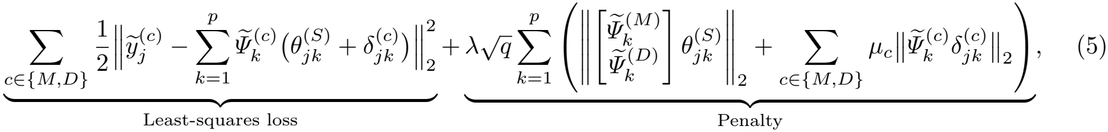

where 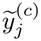 and 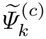 are the residuals of 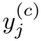 and of 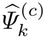 after projecting out the unpenalized columns of cell *c*, the constant and the linear time trend of (2) (Appendix B). Now, we let *A*^(*S*)^, *A*^(*M*)^, *A*^(*D*)^ ∈ R^*p×p*^ denote the adjacency matrices of the scaffold, mother, and daughter networks, respectively. Then, the edges of the network estimates are obtained as 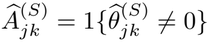 for the scaffold and 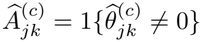 for each cell.

Our estimator requires specifying the order (*q*) of the polynomial basis expansion to approximate the ODE, the tuning parameter for smoothing the observed trajectories to compute 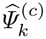 for each cell *c* and protein *k*, and the parameters *λ, µ*_*M*_, and *µ*_*D*_ for the penalty terms of the loss function. We fix *q* in advance based on the setting and the per-cell penalty scaling parameters *µ*_*M*_ = *µ*_*D*_ = 1 so that both network deviations are penalized on the same scale. Then, the smoothing parameter and *λ* are selected jointly through use of the Bayesian information criterion (BIC) [28]. See Appendix B for more details.

### 3.2 Quantifying inheritance

We measure inheritance by how similarly the two estimated networks impact a cell’s protein levels over one sampling interval. We describe the inheritance measure when applied to a PRNODE fit of *q* = 1, and defer the definition of the “true” inheritance, generalizations for *q >* 1 and mathematical properties to Appendix B. Let *J*^(*c*)^ ∈ R^*p×p*^ denote the matrix of the coefficients 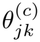 estimated by PRNODE. Then, the fitted ODE model (1) reads 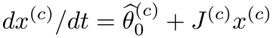. Note that *J*^(*c*)^ when *q* = 1 is simply the Jacobian of the ODE system, which is the same at every state *x*^(*c*)^(*t*). Then, we define *S* = diag (*s*_1_, *s*_2_, …, *s*_*p*_) ∈ R^*p×p*^ as a scaling matrix with the standard deviation of trajectory values for each protein across both cells. Finally, we define 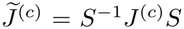 as the scaled Jacobian, capturing the change across proteins at a comparable scale. Over one sampling interval *h*, the regulatory part of this system moves a scaled state *S*^*−*1^*x* ∈ R^*p*^ at time *t* to approximately 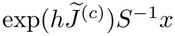 at time *t* + *h*, so we refer to 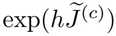 as the “one-step flow map.” Our inheritance measure compares the two cells’ networks through these flow maps. Specifically, we estimate the inheritance between the mother and daughter cells as,

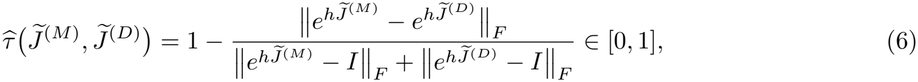

which equals 1 when the two networks coincide. However, as we derive in Appendix E, an inheritance of *τ* = 0 does not refer to the daughter not inheriting regulatory dynamics from her mother; instead, it refers to the daughter inheriting the exact opposite regulatory dynamics (i.e., inhibitors become activators). Instead, an inheritance of *τ ≈* 0.3 refers to no inheritance. See Appendix B for more discussion.

## 4 Simulations

We conduct a simulation study to assess how well PRNODE recovers daughter networks and estimates inheritance when the daughter has fewer observed timepoints than the mother. We seek to understand: 1) how PRNODE compares against competing methods as the daughter trajectory gets shorter, 2) how our tuning procedure compares against plausible alternatives, and 3) how estimation of the network influences the estimation of the inheritance. While we overview particular simulation results in this section, more details and results are provided in Appendix F.

### 4.1 Setup

We consider three regimes: monomial ODE with low noise, monomial ODE with high noise, and cubic ODE with low noise (Figure 3B). For all regimes each cell has 6 proteins, the mother’s time series length is fixed at *n*_*M*_ = 48 (reflecting our real data) and we vary the daughter time series lengths as *n*_*D*_ ∈ *{*12, 24, 36} and the level of inheritance between mother and daughter across four levels (low, mild, moderate, or high; Figure 3A). We draw 20 distinct mother networks as replicates for each regime. This gives 3 *×* 20 *×* 12 = 720 paired mother-daughter time series. We compare PRNODE with GRADE [8], weak SINDy [24], and the kernel ODE (KODE) [11] with a linear kernel. We assess each method’s network recovery by the Matthews correlation coefficient (MCC) [22] over the 30 off-diagonal edges of each cell. Unlike the F1 score, MCC accounts for true negatives and stays informative when most possible edges are absent [9], which is the case in our setting.

**Figure 3.**
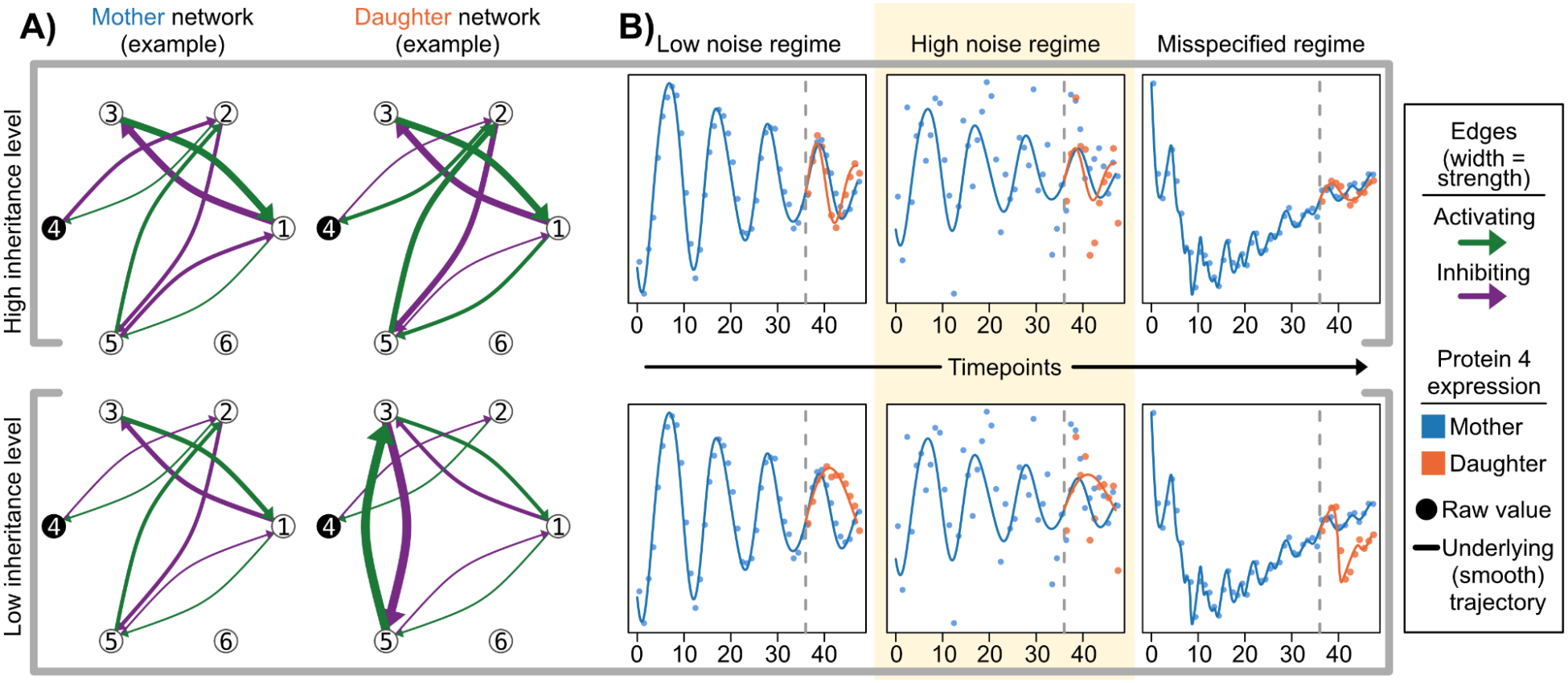
Simulated networks and trajectories. **A**) Simulated network of two mother-daughter pairs at different inheritance levels, where the color and width of each edge depicts the sign and magnitude of the coefficient *θ*_*jk*_, respectively. **B**) Using Protein 4 as an example, the true trajectories as well as observed values are shown for the three different model settings. The dashed vertical line indicates the daughter’s birth time.

### 4.2 PRNODE estimates networks more accurately for shorter daughters than competing methods

When the daughter has a short trajectory (*n*_*D*_ = 12), PRNODE recovers the daughter’s network better than all other methods across every regime and inheritance level (Figure 4). PRNODE also matches or outperforms all other methods across the other regimes, inheritance levels, and daughter lengths, except that KODE outperforms it in the misspecified regime for *n*_*D*_ ∈ {24, 36} at all but the low inheritance level. KODE’s kernel basis plausibly fits the misspecified model more robustly once it has enough daughter observations. Overall, joint estimation improves the estimation of daughter networks over existing methods.

**Figure 4.**
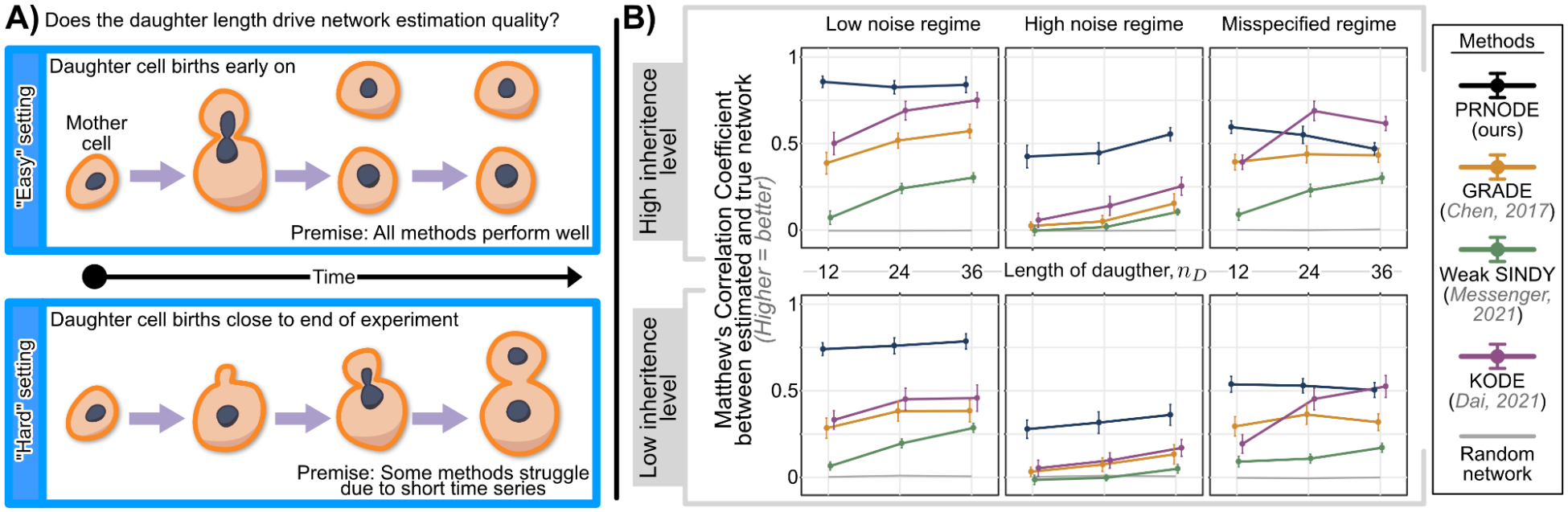
Simulation results of network recovery. **A**) Our premise, where we wish to investigate if daughter cells with shorter trajectories (i.e., born later) yield more difficult estimation tasks, and if PRNODE can overcome this difficulty. **B**) MCC of the estimated daughter network, where higher is better. Columns are the three regimes, and rows are the highest inheritance level (mean true inheritance 0.88) and the lowest (mean true inheritance 0.41). Each method’s mean daughter network MCC with one standard error is shown, and the gray line is the MCC of a random network with the same number of edges as the true daughter network.

### 4.2 Proposed tuning procedure selects tuning parameters appropriately

Because PRNODE’s results rely on the smoothing parameter and the penalty *λ*, we also investigate whether our tuning procedure selects them well. We compare it with two alternatives, (1) fixing the smoothing parameter and selecting only *λ* by BIC and (2) choosing the smoothing parameter and *λ* by cross-validation (CV), on the simulated high-inheritance pairs with *n*_*D*_ = 12 across all regimes (Figure 5A). Our procedure performs better than both alternatives in the low and high noise regimes, and in the misspecified regime it performs similarly to CV and better than the fixed smoothing parameter. We also compare PRNODE’s network recovery at the selected tuning parameters to the best parameters in the search space (“oracle” parameters) across all regimes. The selected networks are not much worse than the oracle networks in the low and high noise regimes, but noticeably worse in the misspecified regime (see Appendix F).

**Figure 5.**
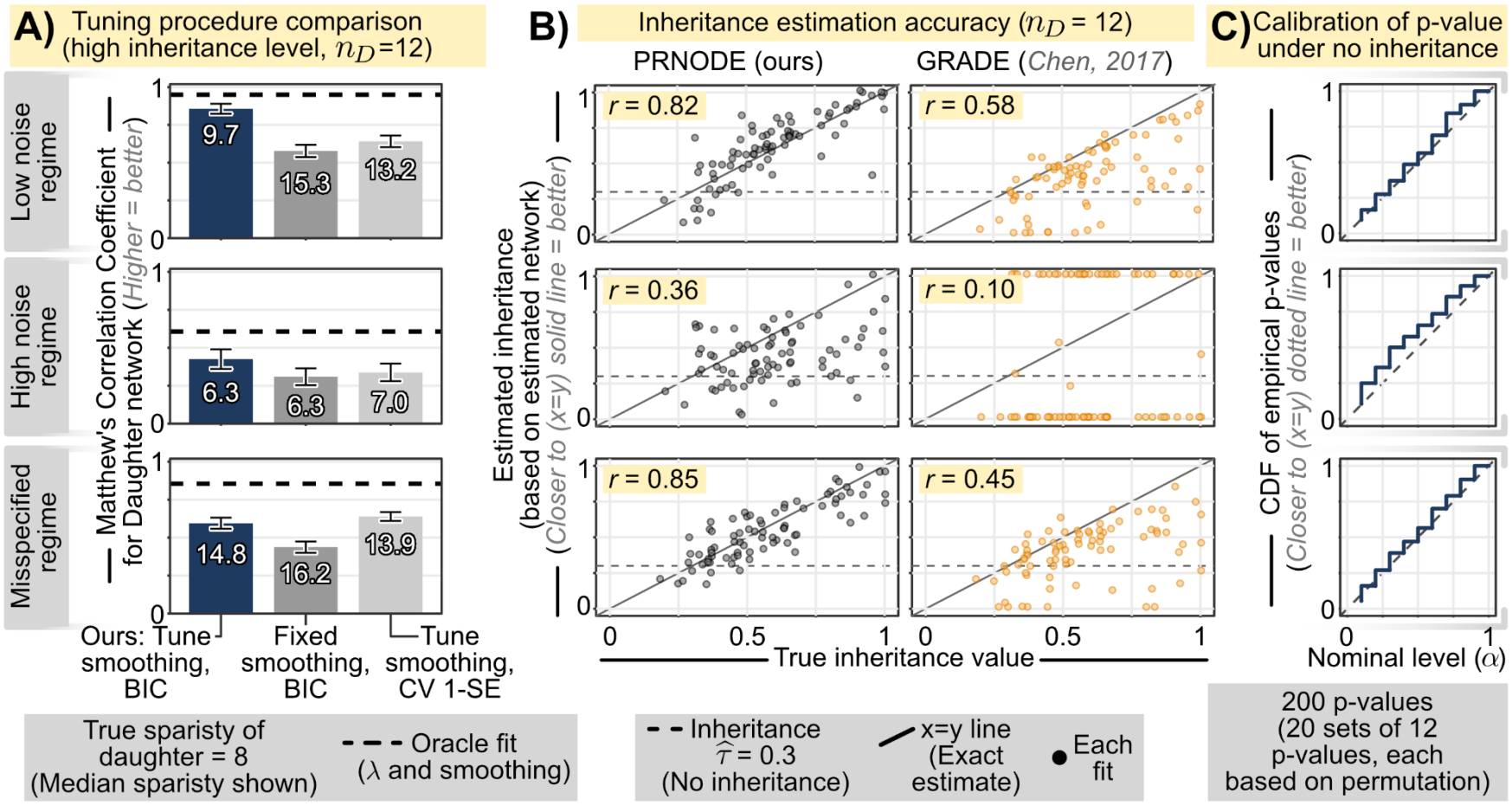
Comparison of tuning procedure and inheritance estimation. **A**) Comparison of different tuning procedures, showing mean and standard error bars, of three different tuning procedures when compared to the oracle fit (dotted), based on the MCC and the mean number of edges of the daughter network. **B**) Estimation of inheritance *τ* when comparing PRNODE and GRADE. **C**) Calibration of estimated inheritance *τ* when computing the empirical p-value comparing the PRNODE fit’s estimate of a daughter and its mother (simulated with no inheritance) and its fit’s estimate of daughter and unrelated mothers.

### 4.4 PRNODE estimates inheritance accurately

We finally assess the quality of the inheritance estimates produced by PRNODE’s estimated networks. We compare the inheritance estimates from PRNODE with those from GRADE, since our inheritance estimate is not straightforward to compute from the networks produced by weak SINDy and KODE. PRNODE’s inheritance estimates correlate better with the true inheritance across all regimes than GRADE’s (Figure 5B). PRNODE’s estimates are noticeably better in the low noise and misspecified regimes than in the high noise regime, which we attribute to the lower variability of the estimates in the two low-noise regimes. We also investigate whether the p-values produced by permutation testing for PRNODE’s inheritance estimate are uniform under a null model. To this end, we generate 20 cohorts of 10 unrelated mother-daughter pairs for each regime with *n*_*D*_ = 12, and run a permutation test for each daughter in a cohort by estimating her network with her true no-inheritance mother and with the 9 unrelated mothers in the same cohort. The p-value for each daughter is the quantile of her inheritance estimate from the fit with her true mother among all PRNODE fits computed for her. Since the true pair has no inheritance, a well-calibrated test gives a uniform distribution of p-values. Figure 5C plots the empirical CDF (ECDF) of the p-values of all 200 pairs for each regime. The p-values are slightly deflated in the high noise regime and slightly inflated in the misspecified regime, so the test is slightly anti-conservative for noisier data and slightly conservative when the model is misspecified, whereas in the low noise regime the ECDF tracks the identity line closely and the test is well calibrated.

## 5 Application to live-cell imaging of budding yeast

Live-cell imaging of budding yeast follows 26 founder cells and their descendants, 207 cells in all, every 12 minutes for 9.4 hours, so that a cell has at most 48 timepoints and a daughter born late in the movie has few (Figure 6A). Six fluorescently tagged proteins are measured in every cell, three that report the position in the division cycle and three that report the response to the environment: Whi5, the G1/S transcriptional repressor whose nuclear exit marks commitment to division; Clb2, the key mitotic cyclin that peaks before anaphase; Cdc10, a septin of the bud-neck ring that assembles after commitment to cell division at the G1/S transition; and Xbp1, Stb3, and Tup1, transcriptional repressors whose nuclear levels respond to stress and starvation rather than to cell-cycle position [3,7]. A FIEST-enabled cell tracking pipeline [20] segments and tracks the cells, records each bud’s birth timepoint and mother, and extracts the protein abundances, giving 180 mother-daughter pairs with a median of 21 daughter timepoints (Figure 6B). The question these data pose is what a dividing cell transmits to its daughter in terms of regulatory dynamics: beyond the protein levels present at birth, does the daughter’s network model her proteins’ trajectories in the same way her mother’s networks models hers?

**Figure 6.**
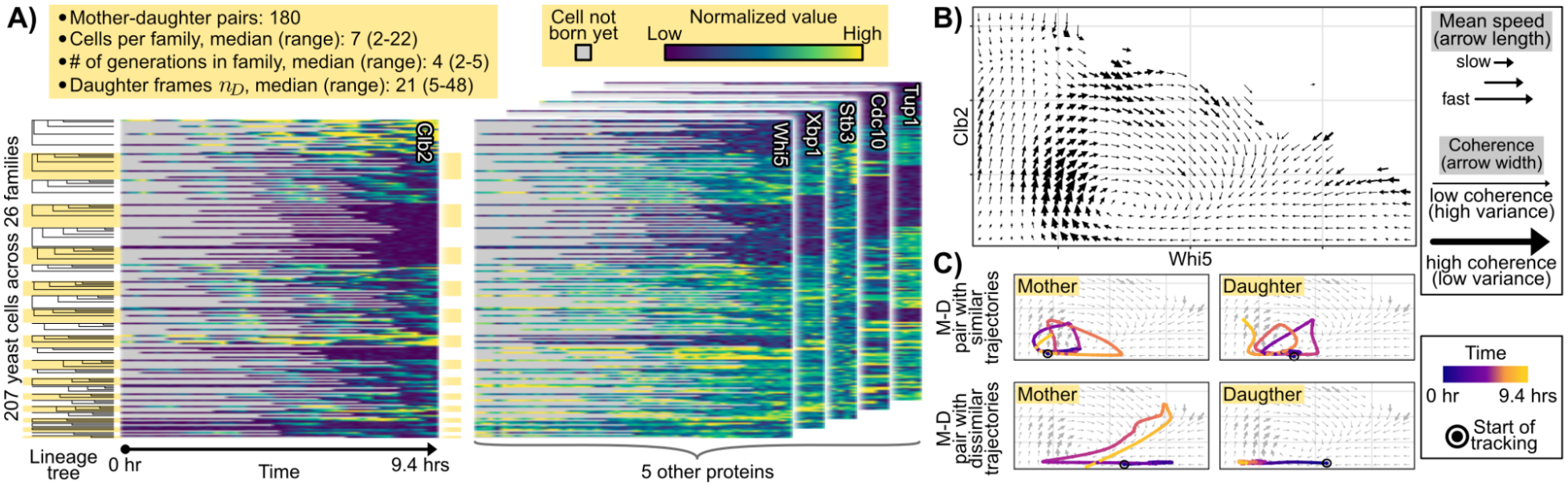
The live-cell data. **A**) Summary of the data, with one heatmap per protein, with the 207 cells as rows, blocked by family and ordered by birth, and the 48 timepoints as columns, gray before the cell’s birth. **B**) Velocity field visualizing where a cell at a given (Whi5, Clb2) value is one timepoint later, averaged over all 207 cells, with arrow length for the mean speed and arrow width for the agreement among cells on the direction. **C**) The smoothed Whi5-by-Clb2 paths of two mother-daughter pairs over the field of (B), colored by the movie’s clock: one pair with similar dynamics and one with dissimilar dynamics.

### 5.1 Mother-daughter pairs move differently through a shared division cycle

To see how varied the dynamics are before any model is fit, we focus on Whi5 and Clb2, which act in opposite phases of the cycle: nuclear Whi5 holds the cell in G1 until Cdk1 activity drives it out, Clb2 then accumulates through mitosis, and its degradation at mitotic exit lets Whi5 return [33,1,7,20]. Figure 6B plots where a cell at each (Whi5, Clb2) position is on average one timepoint later, pooled over all 207 cells, with the arrow’s length for the mean speed and its width for the agreement among cells on the direction. The population traces one counter-clockwise cycle: Clb2 rises at low Whi5, the fastest and most coherent part of the plane, then Whi5 rises as Clb2 falls, and Whi5 falls back at low Clb2. Individual pairs move through this common field very differently (Figure 6C): in one pair the mother and her daughter each complete three loops of the same shape, whereas in another the mother makes one large excursion to high Clb2 while her daughter never leaves low Clb2 in 48 timepoints, which could be explained biologically by the activation of a cell-cycle checkpoint. PRNODE extends this two-protein reading, a shared relation that individual cells follow to different degrees, to all six proteins with one network per cell.

### 5.2 PRNODE estimates reliable, cell-specific networks

PRNODE estimates a different network for each cell, one that reflects the cell’s own trajectory rather than a population-wide model. We fit PRNODE to each of the 180 pairs with the quadratic basis *q* = 2, *µ*_*M*_ = *µ*_*D*_ = 1, giving a median of 12 edges in the mother and 9 in the daughter, 6 of them shared. Figure 7A shows two pairs, one whose mother and daughter are estimated to have similar networks and one whose networks differ. To display the diversity of the networks, we repurpose the flow-map similarity (6) as a dissimilarity between any two estimated networks and embed the 207 cells by multidimensional scaling (Figure 7B; Appendix C). To quantify this heterogeneity, we use “global” GRADE to fit one network to all 207 cells [8], where cells differ only in their initial conditions. The mean squared error (MSE) of this global fit is 0.37 in units of each protein’s population variance, compared to 0.20 for PRNODE’s per-pair fits, and it is worse on every one of the 207 cells (Figure 7C).

**Figure 7.**
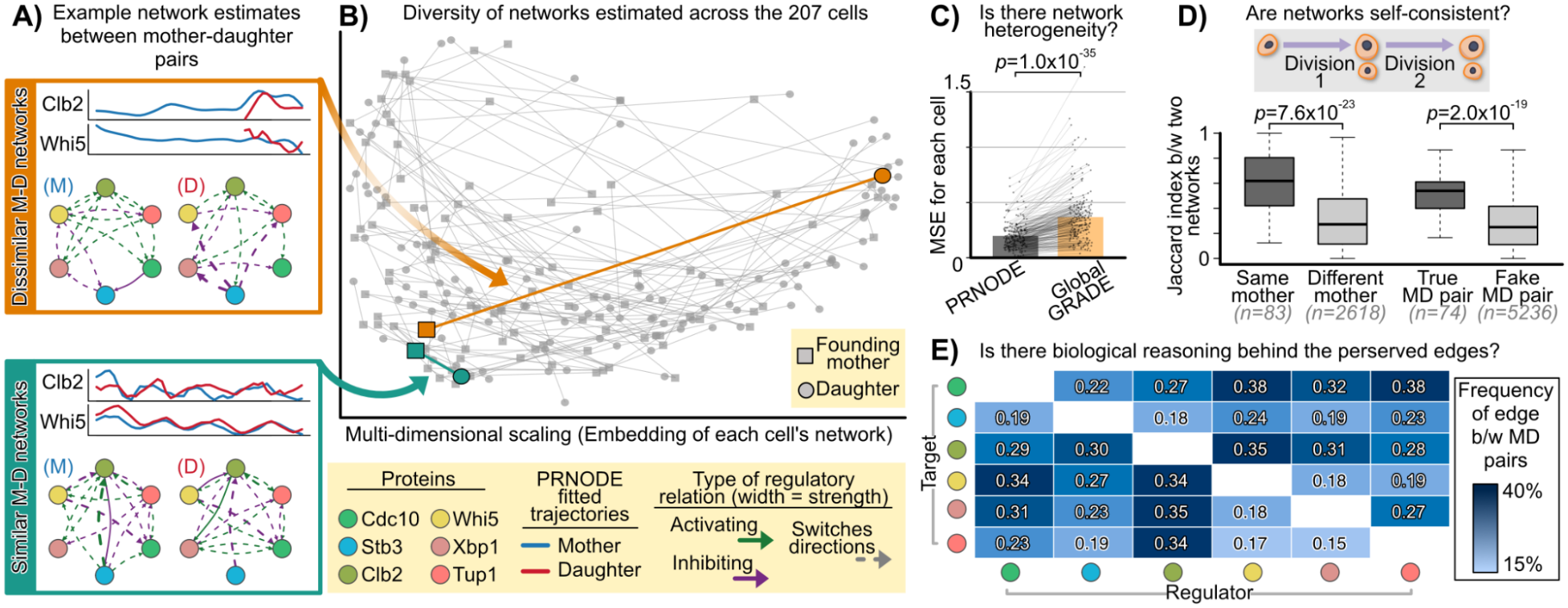
The estimated networks. **A**) Two mother-daughter pairs with similar and with dissimilar estimated networks, showing the fitted PRNODE trajectories for Clb2 and Whi5 and the estimated networks. **B**) Multidimensional scaling of all 207 cells’ networks after applying PRNODE to all 180 pairs of cells. **C**) MSE of each cell when comparing the PRNODE fit to GRADE applied to all the cells simultaneously. Thin lines connect the MSE of each cell in both fits (*p*: Wilcoxon signed-rank test). **D**) Jaccard similarity between the two networks of a mother fitted with two of her daughters, against two different mothers, and between a true mother-daughter pair, against a daughter and an unrelated mother (*p*: Wilcoxon rank-sum test). **E**) Frequency of edges estimated to be shared between both the mother and daughter cell, across the 180 pairs of cells.

Both computational and biological checks support the quality of our networks. Computationally, since a mother of several daughters is fitted once per daughter with a different partner each time, we presume networks driven by noise would disagree between fits of the same mother. Instead, our estimated networks agree at a median Jaccard similarity of 0.62, against 0.27 for the networks of two different mothers, and a daughter’s network agrees with her own mother’s at 0.54, against 0.25 for a daughter and an unrelated mother (Figure 7D; rank-sum *p <* 10^*−*18^ for both). This result demonstrates the self-consistency of the estimated networks. Biologically, the edges most often present in both networks of a pair (Figure 7E) are implicated in the division cycle. For instance, Whi5 inhibits Clb2 and Clb2 activates Whi5 (in 35% and 34% of pairs), the mutual loop of opposite sign behind the cycle of Figure 6B [33,1,7]. Whi5 inhibits the septin Cdc10 (38%), whose ring assembles only after Whi5 has left the nucleus [23], and Clb2 activates the stress repressor Xbp1 (35%), which is transcribed in G2/M [21] and has been shown to be a Cdk1 target with several direct phosphorylation sites including Ser146, Ser271, and Ser551 [34], and the corepressor Tup1 (34%). Appendix G discusses the biology of these edges.

### 5.3 PRNODE detects inheritance across the population more than in individual pairs

We now estimate the inheritance (6) of all 180 pairs (Figure 8A; Appendix C). Because two unrelated networks score about 0.3 rather than 0 (Proposition 4), we read each estimate against a null distribution of 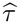 from refitting PRNODE to daughters paired with unrelated mothers, which has a median of 0.26 and a central 95% interval from 0.04 to 0.57. Only 16 of the 180 pairs lie above this centered 95% interval, against the 5 expected by chance (0.025 180 *× ≈* 5). This null asks a strict question: since every cell is a dividing yeast cell, the edges a daughter shares with her mother are largely the consensus edges of Figure 7E, so an unrelated mother is often as similar to a daughter as her own mother is. What the null detects is whether a daughter carries her mother’s deviations from that shared program, much as genetic inheritance is read from the variants a parent passes on rather than from the 99% of the genome common to everyone.

**Figure 8.**
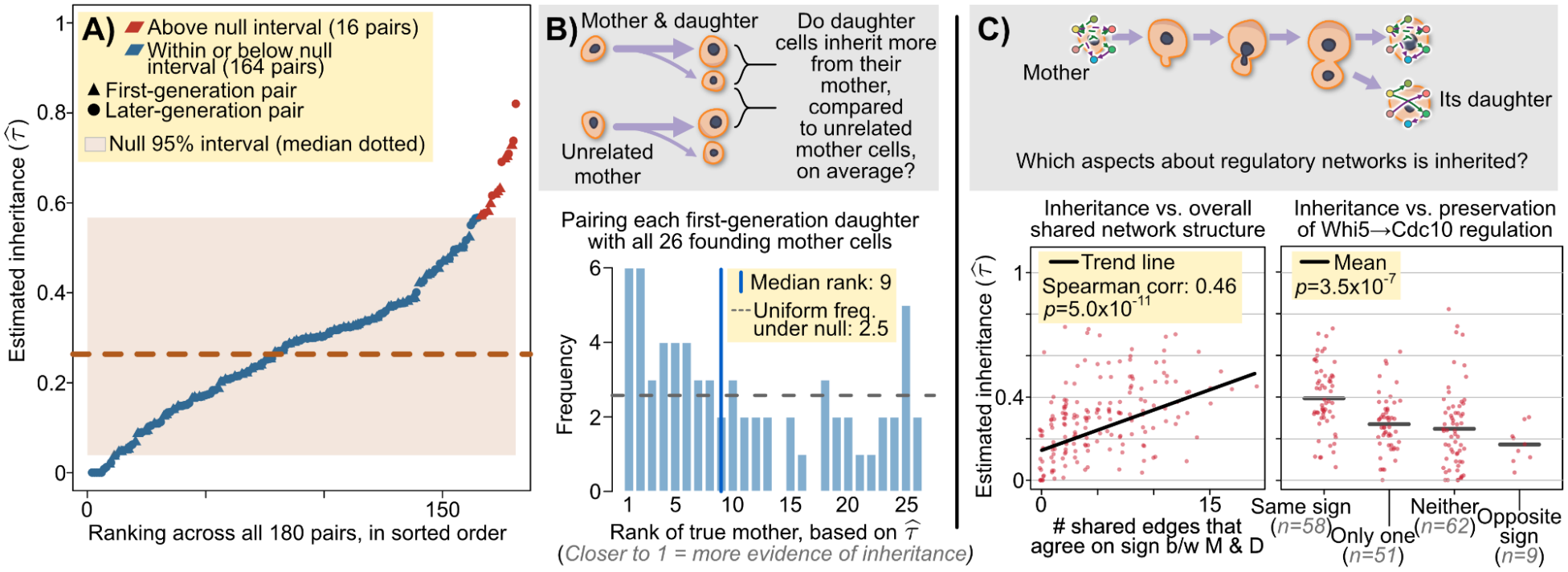
Inheritance across 180 pairs. **A**) The estimated inheritance *τ* across all pairs, with the centered 95% (null) band formed by estimated inheritance when applying PRNODE to daughter-and-unrelated-mother pairs. Dashed line denotes empirical null median. **B**) Rank of the true mother among all 26 founding mothers when PRNODE is applied between each first-generation daughter and every founding mother; rank 1 means no unrelated founder reads as more similar to the daughter than her own mother, and the dashed line is the mean rank under no inheritance. **C**) Estimated inheritance *τ* against the number of shared same-sign edges, and stratified by the sign state of the Whi5 to Cdc10 edge in the two cells (signs evaluated at each cell’s mean state).

Lastly, we show that inheritance is easier to detect across the population than in any one pair. For each of the 67 first-generation daughters we rank her true mother among the 26 founders by the estimated inheritance from 26 separate PRNODE fits, with rank 1 when no unrelated founder reads as more similar than the true mother (Figure 8B). Under no inheritance the rank is uniform with mean 13.5, whereas the observed ranks have a mean of 11.1 and a median of 9, the true mother is in the upper half for 42 of the 67 daughters (binomial *p* = 0.02), and 6 daughters rank their mother first, the smallest attainable level of 1*/*26 for one daughter. The structure that tracks the estimate is the number of shared edges whose direction agrees between mother and daughter (Spearman 0.46, *p* = 5 *×* 10^*−*11^; Figure 8C). Single edges track it weakly: among pairs in which Whi5 inhibits Cdc10 in both cells, the median inheritance is 0.39, against 0.27 when one cell has the edge, 0.25 when neither does, and 0.17 in the 9 pairs where the edge has opposite signs, and although the Kruskal-Wallis test is significant, all four medians lie inside the null interval, so we do not over-interpret the difference (Appendix G).

## 6 Discussion

Live-cell imaging now follows the regulatory proteins of a dividing cell and of her daughters, which raises the question of what a cell transmits to its daughter beyond the proteins present at birth. We frame that question as the similarity between the mother’s and the daughter’s protein regulatory networks, and PRNODE estimates both from one lineage by writing each network as a shared scaffold plus a cell-specific deviation under a group-lasso penalty. In this way, the mother’s longer trajectory supports the daughter’s estimate while the two networks differ where the data demand it. In simulations, the joint fit recovers the daughter’s network better than competing methods, with the largest gains for the shortest daughters, and we find that PRNODE can reliably estimate the inheritance *τ*. On 180 mother-daughter pairs of budding yeast, PRNODE estimates networks that are self-consistent and whose most frequent shared edges recapitulate the division cycle, and we find that the number of shared edges with the same sign tracks our inheritance measure more than any particular regulatory edge.

Our development of PRNODE and analysis of budding yeast motivate us to consider three potential future directions. First, PRNODE fits one pair of cells at a time, so a mother of several daughters is fitted once per daughter and a granddaughter is never fitted with her grandmother. A penalty that follows the full lineage tree, with one deviation per branch, would estimate every network of a family at once and could ask whether a deviation persists across generations. Second, the biological question is often whether a cell’s deviation from a population network, such as the global fit of Figure 7C, is itself inherited, and testing whether a mother’s deviation predicts her daughter’s is the natural next analysis. This could be framed as a hypothesis test that we wish to develop in future work. Third, the inheritance measure takes a value near 0.3 instead of 0 for two unrelated networks (see Appendix E for our theoretical derivations). This can be slightly detrimental for the interpretation of our estimated inheritance. A measure centered at zero under no inheritance, with negative values for opposing dynamics, would be interpretable without an empirical null.

## References

1. Amon, A., Tyers, M., Futcher, B., Nasmyth, K.: Mechanisms that help the yeast cell cycle clock tick: G2 cyclins transcriptionally activate G2 cyclins and repress G1 cyclins. Cell 74(6), 993–1007 (1993)

2. Argüello-Miranda, O., Liu, Y., Wood, N.E., Kositangool, P., Doncic, A.: Integration of multiple metabolic signals determines cell fate prior to commitment. Molecular Cell 71(5), 733–744 (2018)

3. Argüello-Miranda, O., Marchand, A.J., Kennedy, T., Russo, M.A., Noh, J.: Cell cycle–independent integration of stress signals by Xbp1 promotes non-G1/G0 quiescence entry. Journal of Cell Biology 221(1), e202103171 (2021)

4. Bar-Joseph, Z., Gitter, A., Simon, I.: Studying and modelling dynamic biological processes using time-series gene expression data. Nature Reviews Genetics 13(8), 552–564 (2012)

5. Brunton, S.L., Proctor, J.L., Kutz, J.N.: Discovering governing equations from data by sparse identification of nonlinear dynamical systems. Proceedings of the National Academy of Sciences 113(15), 3932–3937 (2016)

6. Cao, J., Zhao, H.: Estimating dynamic models for gene regulation networks. Bioinformatics 24(14), 1619–1624 (2008)

7. Chen, K.C., Calzone, L., Csikasz-Nagy, A., Cross, F.R., Novak, B., Tyson, J.J.: Integrative analysis of cell cycle control in budding yeast. Molecular Biology of the Cell 15(8), 3841–3862 (2004)

8. Chen, S., Shojaie, A., Witten, D.M.: Network reconstruction from high-dimensional ordinary differential equations. Journal of the American Statistical Association 112(520), 1697–1707 (2017). https://doi.org/10.1080/01621459.2016.1229197, 10.1080/01621459.2016.1229197, pMID: 29618851

9. Chicco, D., Jurman, G.: The advantages of the Matthews correlation coefficient (MCC) over F1 score and accuracy in binary classification evaluation. BMC Genomics 21(1), 6 (2020). 10.1186/s12864-019-6413-7

10. Cho, N.H., Cheveralls, K.C., Brunner, A.D., Kim, K., Michaelis, A.C., Raghavan, P., Kobayashi, H., Savy, L., Li, J.Y., Canaj, H., Kim, J.Y.S., Stewart, E.M., Gnann, C., McCarthy, F., Cabrera, J.P., Brunetti, R.M., Chhun, B.B., Dingle, G., Hein, M.Y., Huang, B., Mehta, S.B., Weissman, J.S., Gómez-Sjöberg, R., Itzhak, D.N., Royer, L.A., Mann, M., Leonetti, M.D.: OpenCell: Endogenous tagging for the cartography of human cellular organization. Science 375(6585), eabi6983 (2022)

11. Dai, X., Li, L.: Kernel ordinary differential equations. Journal of the American Statistical Association 117(540), 1711–1725 (2022)

12. Di Talia, S., Wang, H., Skotheim, J.M., Rosebrock, A.P., Futcher, B., Cross, F.R.: Daughter-specific transcription factors regulate cell size control in budding yeast. PLoS Biology 7(10), e1000221 (2009)

13. Gu, Y., Blaauw, D., Welch, J.D.: Bayesian inference of RNA velocity from multi-lineage single-cell data. bioRxiv (2022), doi:10.1101/2022.07.08.499381

14. Herrero, E., Stinus, S., Bellows, E., Berry, L.K., Wood, H., Thorpe, P.H.: Asymmetric transcription factor partitioning during yeast cell division requires the FACT chromatin remodeler and cell cycle progression. Genetics 216(3), 701–716 (2020)

15. Higuchi-Sanabria, R., Pernice, W.M., Vevea, J.D., Alessi Wolken, D.M., Boldogh, I.R., Pon, L.A.: Role of asymmetric cell division in lifespan control in Saccharomyces cerevisiae. FEMS Yeast Research 14(8), 1133–1146 (2014)

16. Hughes, F.A., Barr, A.R., Thomas, P.: Patterns of interdivision time correlations reveal hidden cell cycle factors. eLife 11, e80927 (2022)

17. Huguet, G., Magruder, D.S., Tong, A., Fasina, O., Kuchroo, M., Wolf, G., Krishnaswamy, S.: Mani-fold interpolating optimal-transport flows for trajectory inference. In: Advances in Neural Information Processing Systems. vol. 35 (2022)

18. Ivanov, I.E., Hirata-Miyasaki, E., Chandler, T., Cheloor-Kovilakam, R., Liu, Z., Pradeep, S., Liu, C., Bhave, M., Khadka, S., Arias, C., Leonetti, M.D., Huang, B., Mehta, S.B.: Mantis: high-throughput 4D imaging and analysis of the molecular and physical architecture of cells. PNAS Nexus 3(9), pgae323 (2024)

19. Jindal, K., Adil, M.T., Yamaguchi, N., Yang, X., Wang, H.C., Kamimoto, K., Rivera-Gonzalez, G.C., Morris, S.A.: Single-cell lineage capture across genomic modalities with CellTag-multi reveals fate-specific gene regulatory changes. Nature Biotechnology 42, 946–959 (2024)

20. Kennedy, T., Yalcinkaya, B., Ramakanth, S., Neupane, S., Tadić, N., Buchler, N.E., Argüello-Miranda, O.: Deep learning–driven imaging of cell division and cell growth across an entire eukaryotic life cycle. Molecular Biology of the Cell 36(6), ar74 (2025)

21. Mai, B., Breeden, L.: Xbp1, a stress-induced transcriptional repressor of the Saccharomyces cerevisiae Swi4/Mbp1 family. Molecular and Cellular Biology 17(11), 6491–6501 (1997)

22. Matthews, B.W.: Comparison of the predicted and observed secondary structure of T4 phage lysozyme. Biochimica et Biophysica Acta (BBA)-Protein Structure 405(2), 442–451 (1975). 10.1016/0005-2795(75)90109-9

23. McMurray, M.A., Thorner, J.: Septins: molecular partitioning and the generation of cellular asymmetry. Cell Division 4, 18 (2009)

24. Messenger, D.A., Bortz, D.M.: Weak SINDy: Galerkin-based data-driven model selection. Multiscale Modeling & Simulation 19(3), 1474–1497 (2021)

25. Mura, M., Feillet, C., Bertolusso, R., Delaunay, F., Kimmel, M.: Mathematical modelling reveals unexpected inheritance and variability patterns of cell cycle parameters in mammalian cells. PLoS Computational Biology 15(6), e1007054 (2019)

26. Reinbold, P.A.K., Gurevich, D.R., Grigoriev, R.O.: Using noisy or incomplete data to discover models of spatiotemporal dynamics. Physical Review E 101(1), 010203 (2020)

27. Sandler, O., Mizrahi, S.P., Weiss, N., Agam, O., Simon, I., Balaban, N.Q.: Lineage correlations of single cell division time as a probe of cell-cycle dynamics. Nature 519(7544), 468–471 (2015)

28. Schwarz, G.: Estimating the dimension of a model. The Annals of Statistics 6(2), 461–464 (1978)

29. Siahpirani, A.F., Knaack, S., Chasman, D., Seirup, M., Sridharan, R., Stewart, R., Thomson, J., Roy, S.: Dynamic regulatory module networks for inference of cell type-specific transcriptional networks. Genome Research 32(7), 1367–1384 (2022)

30. Simon, N., Tibshirani, R.: Standardization and the group lasso penalty. Statistica Sinica 22(3), 983–1001 (2012), http://www.jstor.org/stable/24309971

31. Singh, R., Wu, A.P., Berger, B.: Granger causal inference on DAGs identifies genomic loci regulating transcription. In: International Conference on Learning Representations (2022)

32. Singh, R., Wu, A.P., Mudide, A., Berger, B.: Causal gene regulatory analysis with RNA velocity reveals an interplay between slow and fast transcription factors. Cell systems 15(5), 462–474 (2024)

33. Skotheim, J.M., Di Talia, S., Siggia, E.D., Cross, F.R.: Positive feedback of G1 cyclins ensures coherent cell cycle entry. Nature 454(7202), 291–296 (2008)

34. Tao, R., Chen, H., Gao, C., Xue, P., Yang, F., Han, J.D.J., Zhou, B., Chen, Y.G.: Xbp1-mediated histone h4 deacetylation contributes to dna double-strand break repair in yeast. Cell Research 21(11), 1619–1633 (Nov 2011). https://doi.org/10.1038/cr.2011.58, 10.1038/cr.2011.58

35. Tyson, J.J., Novák, B.: Models in biology: Lessons from modeling regulation of the eukaryotic cell cycle. BMC Biology 13, 1–10 (2015)

36. Weinreb, C., Rodriguez-Fraticelli, A., Camargo, F.D., Klein, A.M.: Lineage tracing on transcriptional landscapes links state to fate during differentiation. Science 367(6479), eaaw3381 (2020)

37. Welch, J.D., Hartemink, A.J., Prins, J.F.: SLICER: inferring branched, nonlinear cellular trajectories from single cell RNA-seq data. Genome Biology 17, 106 (2016)

38. Wu, H., Lu, T., Xue, H., Liang, H.: Sparse additive ordinary differential equations for dynamic gene regulatory network modeling. Journal of the American Statistical Association 109(506), 700–716 (2014)

39. Wu, W., Kennedy, T., Arguello-Miranda, O., Lin, K.Z.: Measuring regulatory network inheritance in dividing yeast cells using ordinary differential equations. bioRxiv pp. 2024–11 (2025)

40. Yuan, M., Lin, Y.: Model selection and estimation in regression with grouped variables. Journal of the Royal Statistical Society: Series B (Statistical Methodology) 68(1), 49–67 (2006)

41. Zhang, J., Larschan, E., Bigness, J., Singh, R.: scNODE: Generative model for temporal single cell transcriptomic data prediction. Bioinformatics 40(Supplement_2), ii146–ii154 (2024)

42. Zhang, S., Pyne, S., Pietrzak, S., Halberg, S., McCalla, S.G., Siahpirani, A.F., Sridharan, R., Roy, S.: Inference of cell type-specific gene regulatory networks on cell lineages from single cell omic datasets. Nature Communications 14, 3064 (2023)

