## Appendix for "Joint Estimation of Mother and Daughter Protein Regulatory Networks Quantifies Inheritance in Yeast Lineages"

#### Code and data availability

PRNODE is implemented in the R package `LICmethods` (MIT license), whose repository at `https://github.com/shirmath/LIC_methods` carries a README, the package vignette, and the scripts that reproduce every figure and number in this paper, including the simulations of Section 4 and the yeast analyses of Section 5.

#### Author contributions

S.M. and K.Z.L. developed the method, wrote the software, carried out the analyses, and wrote the manuscript. W.W. preprocessed the imaging data and prepared the single-cell trajectories. T.K. and O.A.-M. designed and performed the live-cell imaging experiments and the segmentation and tracking. O.A.-M. guided the biological interpretation. K.Z.L. supervised the project and secured funding. All authors read and approved the manuscript.

#### Funding

This work was supported by the National Institute of General Medical Sciences of the National Institutes of Health [R35GM162089 to K.Z.L. and R00GM135487 to O.A.-M.]; the National Institute for Theory and Mathematics in Biology, funded by the National Science Foundation [DMS-2235451] and the Simons Foundation [MPS-NITMB-00005320]; and the National Science Foundation [DMS-2424748].

#### Acknowledgements

We thank the members of the Argüello-Miranda and Lin laboratories for discussions of the lineage analyses. The authors used Claude (Anthropic) to draft and revise portions of the text and code of this manuscript; the authors reviewed all of its output and take full responsibility for the content.

#### Competing interests

None declared.

#### Organization of the appendix

Appendix A compares PRNODE with the existing frameworks for inferring cellular dynamics from single-cell data. Appendix B gives the details of PRNODE: the stacked group-lasso form of the objective, the block coordinate descent that solves it, the tuning procedure, and the inheritance measure. Appendix C describes the analyses of the main text that follow a PRNODE fit. Appendix D gives the pseudocode of the whole procedure. Appendix E states the theoretical properties of PRNODE that clarify the roles of the tuning parameters and of the inheritance measure, with the proofs in Appendix H. Appendix F gives additional simulation results, and Appendix G additional results on the 180 mother-daughter pairs of yeast cells.

⋆ Corresponding author.

### A Extended background

#### A.1 Single-cell snapshot and lineage-tracing analyses

Most cell-level measurements of gene regulation come from single-cell RNA sequencing, which destroys each cell it measures, so that every cell contributes one snapshot and the order of cells in time must be inferred. Pseudotime methods order cells by the similarity of their transcriptomes, through principal curves fitted along a tree of clusters in Slingshot [27] or through a Markov chain on a neighbor graph in Palantir [23], whose fate probabilities describe the likely trajectories of the population rather than the path of any one cell. RNA velocity estimates the time derivative of each gene's spliced abundance from its ratio of unspliced to spliced reads under a splicing ODE [12,3], and scVelo pools the per-gene fits into one latent time per cell, although the direction of a gene's dynamics is not identifiable when the data capture only part of its trajectory [3]. TFvelo comes closest to a regulatory network by modeling the derivative of a target gene's abundance as a linear function of the abundances of its candidate transcription factors, but it fits that model along an inferred time whose direction is fixed by constraining every degradation rate to be positive [13]. When cells are sampled at several times, neural ODEs such as MIOFlow learn a flow that carries the cells measured at one time onto those measured at the next under an optimal-transport loss [9], a model of how the population moves, since no cell is measured at more than one time. MultistageOT likewise describes the population, ordering the cells of one snapshot into successive stages through a chain of optimal-transport couplings [29].

Lineage tracing supplies the relationships between cells that snapshots lack. Heritable barcodes, read out alongside the transcriptome [4,30] or alongside chromatin accessibility [10], identify which sequenced cells descend from a common ancestor, and they show that fate is heritable during reprogramming, with a correlation of 0.71 between the reprogramming rate of a clone and that of its parent clone [4]. The same labels expose what expression alone misses, because a velocity-based trajectory method applied to the CellTag-multi data does not recover the early states that the clones show to lead to distinct fates [10]. Barcodes are still read only when cells are sequenced, so that a clone records membership at a few timepoints and leaves the dynamics of each cell between those points unobserved.

#### A.2 SINDy and weak SINDy

SINDy [6] writes each $f_j$ in $dx(t)/dt = f(x(t))$ as a linear combination $\sum_{l=1}^{L} \xi_{jl} g_l(x)$ of candidate functions, such as monomials in the components of $x$, and estimates a sparse $\xi_j \in \mathbb{R}^L$ by regressing a numerical derivative of $x_j$ on the candidates. Because a numerical derivative amplifies measurement error, weak formulations multiply the ODE by smooth test functions and integrate by parts, so that the derivative falls on the known test function rather than on the data [21,18]. Reinbold et al. show that this approach estimates coefficients accurately at noise levels at which derivative-based sparse regression does not, and that sparsity removes the extraneous terms of a library larger than the true one, so that the exact form of the right-hand side need not be known in advance [21]. Weak SINDy [18] still needs enough timepoints inside the support of each test function to evaluate its integrals, which a daughter observed a dozen times does not provide.

#### A.3 The GRADE model

GRADE [8] assumes that each $f_j$ is additive in the components of $x$ and approximates each additive component by a linear combination of $q$ basis functions, the model (1) with the polynomial basis that we use (GRADE allows other bases). Because (1) is linear in the coefficients, integrating both sides expresses the trajectory itself, rather than its derivative, as a linear function of the coefficients, which is the model (2). GRADE replaces each $x_k^{(c)}$ by a smoothing-spline estimate $\widehat{x}_k^{(c)}$ whose smoothing parameter is chosen by generalized cross-validation, computes the integrated basis $\widehat{\Psi}_k^{(c)}(t)$, and fits (2) to one cell at a time, with $n_c$

the number of timepoints of cell $c$, by solving, for each protein $j \in \{1, \dots, p\}$,

$$\min_{C_{j0}^{(c)}, \theta_{j0}^{(c)}, \theta_{j1}^{(c)}, \dots, \theta_{jp}^{(c)}} \frac{1}{2n_c} \sum_{i=1}^{n_c} \left\{ y_{ij}^{(c)} - C_{j0}^{(c)} - \theta_{j0}^{(c)} (t_i^{(c)} - t_1^{(c)}) - \sum_{k=1}^{p} \widehat{\Psi}_k^{(c)} (t_i^{(c)})^\top \theta_{jk}^{(c)} \right\}^2 + \lambda \sum_{k=1}^{p} \left[ \frac{1}{n_c} \sum_{i=1}^{n_c} \{ \widehat{\Psi}_k^{(c)} (t_i^{(c)})^\top \theta_{jk}^{(c)} \}^2 \right]^{1/2}, \tag{S1}$$

where $y_{ij}^{(c)}$ is the observed level of protein $j$ at timepoint $t_i^{(c)}$, $C_{j0}^{(c)}$ estimates the initial condition $x_j^{(c)}(t_1^{(c)})$, $\theta_{j0}^{(c)}$ is the constant term of (1), whose integrated column is the linear time trend $t_i^{(c)} - t_1^{(c)}$, and the group lasso [31] penalizes the fitted contribution of each regulator $\theta_{jk}^{(c)} \in \mathbb{R}^q$ [24]. Chen et al. prove that GRADE recovers the true network with probability tending to one as the number of observations grows under a correctly specified model, with the number of proteins allowed to grow as well, and they extend (S1) to several series that share one set of coefficients but differ in their initial conditions [8].

### B Details of the PRNODE estimator

#### B.1 Data scaling, profiling details, and equivalence to group lasso

Here, we provide details on data pre-processing for our method, profiling out unpenalized terms from the loss, and the equivalence of our objective to a group lasso problem. First, we smooth the trajectories for all the proteins in each cell via fitting a cubic smoothing spline with a knot at every timepoint to each protein's observed time series. Then, both the observed values and smoothed trajectories of each protein are divided by the pooled standard deviation of its observed values, denoted $s_j$, computed over the observations of both cells. This ensures that every protein is on a comparable scale prior to fitting PRNODE. Then, we compute the integrated basis functions $\widehat{\Psi}_k^{(c)}(t_i^{(c)})$ for all proteins and cells in (2) from the scaled smoothed series by the trapezoidal rule on a grid of 2,000 steps per sampling interval.

We then profile out each cell's initial condition and constant term to obtain $\widetilde{y}_j^{(c)}$ and $\widetilde{\Psi}^{(c)}$ as in (5). Let $n_D, n_M$ denote the number of timepoints observed for the daughter and mother, respectively, and define $Z^{(c)} = \begin{bmatrix} \mathbf{1} \; t_i^{(c)} \end{bmatrix} \in \mathbb{R}^{n_c \times 2}$. We obtain closed-form solutions for the minimizers of the loss in $C_{j0}^{(c)}$ and $\theta_{j0}^{(c)}$ as

$$\begin{bmatrix} \widehat{C}_{j0}^{(c)} \\ \widehat{\theta}_{j0}^{(c)} \end{bmatrix} = \left( Z^{(c)\top} Z^{(c)} \right)^{-1} Z^{(c)\top} \left( y_j^{(c)} - \sum_{k=1}^{p} \widehat{\Psi}_k^{(c)} \left( \theta_{jk}^{(S)} + \delta_{jk}^{(c)} \right) \right)$$

Plugging the above closed-form estimates back into the loss yields the profiled loss of (5), with

$$\widetilde{y}_j^{(c)} = (I - H_c) y_j^{(c)} \qquad \text{and} \qquad \widetilde{\Psi}_k^{(c)} = (I - H_c) \widehat{\Psi}_k^{(c)},$$

where

$$H_c = Z^{(c)} \left( Z^{(c)\top} Z^{(c)} \right)^{-1} Z^{(c)\top}$$

is the projection onto the two unpenalized columns of cell $c$.

Now, we describe how our profiled loss can be equivalently stated as a group lasso problem. We collect the profiled responses of the two cells as $\widetilde{\mathbf{y}}_j = \left[ \widetilde{y}_j^{(M)\top}, \; \widetilde{y}_j^{(D)\top} \right]^\top \in \mathbb{R}^{n_M + n_D}$, and build the design matrix

$$\widetilde{\boldsymbol{\Psi}} = \begin{bmatrix} \widetilde{\Psi}^{(M)} & \widetilde{\Psi}^{(M)} & 0 \\ \widetilde{\Psi}^{(D)} & 0 & \widetilde{\Psi}^{(D)} \end{bmatrix} \in \mathbb{R}^{(n_M + n_D) \times 3pq}, \tag{S2}$$

whose $3p$ column groups, each with $q$ columns, are $X_{S,k}$ (the scaffold block, with $X_{S,k} = X_{M,k} + X_{D,k}$), $X_{M,k}$, and $X_{D,k}$ for $k \in \{1, \dots, p\}$, where $X_{M,k}$ and $X_{D,k}$ hold $\widetilde{\Psi}_k^{(c)}$ in the rows of one cell and zeros in the other's. We also visualize this design in Figure S1 to map out which columns are used in estimation of which coefficients.

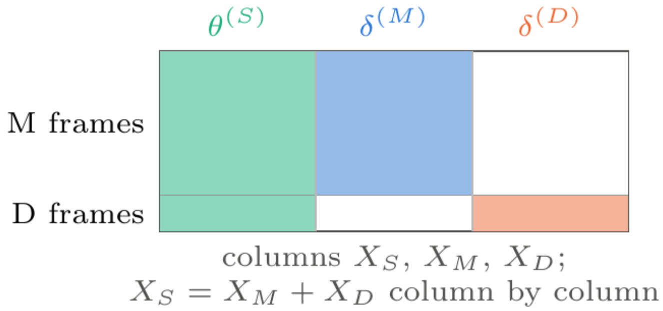


**Figure S1.** Diagram of how design matrix for group lasso problem is constructed.

Writing $\theta^{(S)}_{jk}$, $\delta^{(M)}_{jk}$, and $\delta^{(D)}_{jk}$ for the coefficients of these groups, the profiled objective (5) becomes

$$\min_{\boldsymbol{\theta}_j \in \mathbb{R}^{3pq}} \ \frac{1}{2}\|\tilde{\mathbf{y}}_j - \widetilde{\boldsymbol{\Psi}}\boldsymbol{\theta}_j\|_2^2 + \lambda\sqrt{q}\sum_{k=1}^{p}\Big(\|X_{S,k}\theta^{(S)}_{jk}\|_2 + \mu_M\|X_{M,k}\delta^{(M)}_{jk}\|_2 + \mu_D\|X_{D,k}\delta^{(D)}_{jk}\|_2\Big), \tag{S3}$$

where $\boldsymbol{\theta}_j$ stacks the $3p$ coefficient groups and no column is unpenalized. This is the standardized group lasso of Simon and Tibshirani [24] for the design $\widetilde{\boldsymbol{\Psi}}$ and the $3p$ groups, with the group weights $\sqrt{q}$, $\sqrt{q}\mu_M$, and $\sqrt{q}\mu_D$.

##### B.2 Objective function properties and optimization

The objective (S3) is convex in its parameters because it is a group lasso problem, and it is separable across the $p$ proteins, so PRNODE is $p$ group-lasso problems that share one $\lambda$ and the same design. We minimize the objective using cyclic block coordinate descent, the standard approach for group-lasso problems [31,17]. Each group's columns are orthonormalized by a thin QR decomposition, $X_{g,k} = Q_{g,k}R_{g,k}$, so that in the coordinates $b_{g,k} = R_{g,k}\beta_{g,k}$ the penalty on the group is $\lambda\sqrt{q}\mu_g\|b_{g,k}\|_2$ and the exact minimizer over one group, with the others held fixed, is the group soft-thresholding step $b_{g,k} \leftarrow \big(1 - \lambda\sqrt{q}\mu_g/\|Q^\top_{g,k}r_{g,k}\|_2\big)_+ Q^\top_{g,k}r_{g,k}$, where $r_{g,k}$ is the residual of $\tilde{\mathbf{y}}_j$ on all other groups. The solver cycles these steps over the groups, warm-starting along a decreasing path of $\lambda$ from $\lambda_{\max}$ of Proposition 1, screens groups with the sequential strong rule [28], and stops at each $\lambda$ when the duality gap [5] and the Karush-Kuhn-Tucker conditions are within a relative tolerance of $10^{-10}$. The estimates are read back as $\widehat{\beta}_{g,k} = R^{-1}_{g,k}\widehat{b}_{g,k}$, and a regulator is declared an edge when the norm of its fitted contribution exceeds $10^{-8}\|\tilde{\mathbf{y}}_j\|_2$. This tolerance allows PRNODE to ignore very small coefficient magnitudes that might arise due to numerical issues.

##### B.3 Tuning procedure

We use BIC to select the smoothing parameter and $\lambda$ jointly. We fit PRNODE for each $\lambda$ and smoothing parameter combination over a grid of candidate values for each parameter. The $\lambda$ grid has 80 values, log-spaced over four decades from $\lambda_{\max}$ down to $10^{-4}\lambda_{\max}$, where $\lambda_{\max}$ is the largest of the $p$ proteins' values of Proposition 1 so that the grid is the same for every protein. The grid for the smoothing parameter is $\{0, 0.25, 0.5, 0.75\}$ on the scale of `spar`. For each value of the smoothing parameter we build the design as described above and fit the full $\lambda$ path. Then, we choose the smoothing parameter and $\lambda$ as the combination that minimizes

$$\mathrm{BIC}(\texttt{spar}, \lambda) = \sum_{j=1}^{p}\Big[\mathrm{RSS}_j + \log(n)\,k_j\Big], \quad \text{where} \quad n = n_M + n_D, \tag{S4}$$

where $k_j$ is the number of groups of the $3p$ with a nonzero coefficient in protein $j$'s fit at $(\texttt{spar}, \lambda)$, and $\mathrm{RSS}_j$ is the residual sum of squares of the weighted responses, written as

$$\mathrm{RSS}_j = \omega \sum_{c\in\{M,D\}} \frac{1}{2}\Big\|\tilde{y}^{(c)}_j - \sum_{k=1}^{p}\widetilde{\Psi}^{(c)}_k\big(\theta^{(S)}_{jk} + \delta^{(c)}_{jk}\big)\Big\|_2^2. \tag{S5}$$

In (S5), $\omega$ is defined as the inverse of the scaled estimated variance of the noise in our observed realizations. Including $\omega$ in the RSS puts the fits of pairs with different numbers of timepoints on a comparable scale. We compute $\omega$ as

$$\omega = \frac{n_M + n_D}{n_M/\omega_M + n_D/\omega_D},$$

where $\omega_c = 1/\left[\frac{1}{p}\sum_{j=1}^{p}(\widehat{\sigma}_{cj}/s_j)^2\right]$ and $\widehat{\sigma}_{cj}$ is the standard deviation of the second differences of the observed (unscaled) series of protein $j$ for cell $c$ divided by $\sqrt{6}$ and serves as an estimate of the noise standard deviation. The form of $\widehat{\sigma}_{cj}$ is a plug-in estimate for the noise standard deviation computed under the assumption that the noise terms are independent across timepoints and the trajectory for each cell and protein is locally linear. Since we use the weighted residual sum-of-squares, the criterion in (S4) is the BIC [22] of a Gaussian model with $1/\omega$ plugged in for the error variance.

As we demonstrate in Appendix E, tuning $\mu_D$ and $\mu_M$ in practice might lead to detrimental side-effects where the $\widehat{\theta}_{jk}^{(S)}$ becomes empty (i.e., no sharing of information) or forces $\widehat{\delta}_j^{(c)} = 0$ for one of the two cells (i.e., "leakage" of information). Hence, we typically fix $\mu_M = \mu_D = 1$ and only vary $\mu_D$ and $\mu_M$ to assess sensitivity.

##### B.4 Details on inheritance measure

We discuss additional nuances to calculate the inheritance metric $\tau$ defined in (6) as well as properties and qualities of this metric. While the main text describes the inheritance metric for $q = 1$, we now describe its more general form when $q > 1$. For $q > 1$ the Jacobian of the polynomial system depends on the state, so we replace $J^{(c)}$ by the Jacobian of the system evaluated at each cell's respective mean-state (i.e., the average protein value for that cell over its trajectory). For $q > 1$ the right-hand side of (1) for cell $c$ is a polynomial field $f^{(c)}$ whose $j$th component is $\theta_{j0}^{(c)} + \sum_{k=1}^{p}\sum_{m=1}^{q}\theta_{jk,m}^{(c)}x_k^m$, with $\theta_{jk,m}^{(c)}$ the $m$th entry of $\theta_{jk}^{(c)}$, and its Jacobian $J^{(c)}(x) \in \mathbb{R}^{p\times p}$, with entries $\partial f_j^{(c)}/\partial x_k = \sum_{m=1}^{q} m\,\theta_{jk,m}^{(c)}x_k^{m-1}$, changes with the state $x$. We linearize each cell's field at its own mean state $\overline{x}^{(c)} \in \mathbb{R}^p$, the mean of the cell's protein levels over its timepoints, and set $J^{(c)} = J^{(c)}(\overline{x}^{(c)})$, which at $q = 1$ is the coefficient matrix of Section 3.2 whatever the state. The two linearized systems are then standardized, $\widetilde{J}^{(c)} = S^{-1}J^{(c)}S$ with $S = \mathrm{diag}(s_1, \ldots, s_p)$ the pooled standard deviations of the two cells' protein levels, so that each protein's rate of change is measured in units of its own variability, and compared through their one-step flow maps $\exp(h\widetilde{J}^{(c)})$ in (6), with $h$ one sampling interval.

The *true inheritance* of a pair applies this calculation to the true coefficients: each cell is linearized at the mean of its own noiseless trajectory, and $S$ holds the standard deviations of the noiseless trajectories pooled over the two cells. It is the target that the simulations of Section 4 plot on the horizontal axis. The *plug-in estimate* $\widehat{\tau}$ replaces each ingredient by its sample counterpart: the PRNODE coefficients $\widehat{\theta}_{jk}^{(c)} = \widehat{\theta}_{jk}^{(S)} + \widehat{\delta}_{jk}^{(c)}$ of the penalized fit at the BIC-selected smoothing parameter and $\lambda$ (not the least-squares refit used inside (S4)), the mean of each cell's observed levels as its state of linearization, and the pooled standard deviations $s_j$ of the observed intensities defined above as $S$. Because PRNODE fits the scaled data, the coefficients returned by the solver are already on this standardized scale, and the estimate exponentiates their Jacobians at the standardized mean states directly.

The measure has certain appealing properties.

1. **Bounded in** $[0, 1]$: $\tau \in [0, 1]$ by the triangle inequality, with $\tau = 1$ if and only if the two networks coincide and $\tau = 0$ if and only if the two one-step displacements point in opposite directions, so that 0 is opposing and 1 is aligned.
2. **Linear approximation**: Because $\Phi^{(c)} - I = h\widetilde{J}^{(c)} + O(h^2)$ in Big-O notation,

$$\tau = 1 - \frac{\|\widetilde{J}^{(M)} - \widetilde{J}^{(D)}\|_F}{\|\widetilde{J}^{(M)}\|_F + \|\widetilde{J}^{(D)}\|_F} + O(h),$$

so the sampling interval cancels at first order and $\tau$ is one minus a normalized distance between the two generators of the dynamics.

3. **Divergence of trajectories**: For any state $x$, $\|(\Phi^{(M)} - \Phi^{(D)})x\|_2 \le (1-\tau)\big(\|\Phi^{(M)} - I\|_F + \|\Phi^{(D)} - I\|_F\big)\|x\|_2$, so $1-\tau$ bounds how far the two cells' one-step predictions from a common state diverge, relative to the size of the moves themselves.
4. **Caveat on what "no inheritance" means**: Two unrelated networks do not have $\tau = 0$: two networks of equal size whose coefficient matrices are orthogonal, $\mathrm{tr}(\widetilde{J}^{(M)\top}\widetilde{J}^{(D)}) = 0$, have $\tau$ near $1 - 1/\sqrt{2} \approx 0.3$ (Proposition 4 and its proof in Appendix H), so an estimate of inheritance should be read against this null value rather than against zero.

Comparing the one-step flow maps rather than the coefficients or the edge sets weights each coefficient by its effect on the trajectory, so that an edge too weak to move the trajectory barely changes $\tau$.

### C Details of downstream methods after fitting PRNODE

#### C.1 Velocity field visualization

Figure 6B shows, for a cell at a given (Whi5, Clb2) value, where the cell is one timepoint later, averaged over all 207 cells. Each cell's Whi5 and Clb2 series are smoothed by the same cubic smoothing spline used before fitting PRNODE, and each cell contributes one transition per consecutive pair of its timepoints, namely its smoothed (Whi5, Clb2) position at a timepoint and its displacement, the smoothed value one timepoint later minus the value at that timepoint, for 5,162 transitions in all. The field is evaluated on a regular 40-by-20 grid over the box from the 1st to the 99th percentile of the positions on each axis. At each gridpoint the transitions are averaged with a Gaussian kernel in position whose standard deviation on each axis is 1.5 grid spacings. The arrow at a gridpoint points along the kernel-weighted mean displacement; its length is a monotone transform of the mean speed, with each axis measured in units of that protein's standard deviation and speeds above an upper quantile capped so that a few fast gridpoints do not shrink the rest; and its width is the coherence of the direction, the mean resultant length $\|\sum_i w_i u_i\|_2 / \sum_i w_i$ of the unit displacement vectors $u_i$ under the kernel weights $w_i$, which is 1 when every nearby transition points the same way and near 0 when they cancel. Gridpoints supported by too few transitions are left blank. Figure 6C overlays the smoothed paths of two mother-daughter pairs on the same field, colored by time since the start of the movie.

#### C.2 Network visualization

A network is drawn with the $p = 6$ proteins as nodes and an arrow from regulator $k$ to protein $j$ whenever the estimated group $\widehat{\theta}^{(c)}_{jk} \in \mathbb{R}^q$ of cell $c$ is nonzero (Figure 7A). At $q = 1$ the group is a single coefficient, whose sign gives the direction of the edge, activating when positive and inhibiting when negative, and whose magnitude gives its strength. At $q > 1$ the contribution of regulator $k$ to $dx_j/dt$ is the polynomial $\theta^{(c)}_{jk1}x_k + \cdots + \theta^{(c)}_{jkq}x_k^q$, whose effect on protein $j$ depends on the current level of protein $k$, so we read both the direction and the strength from the derivative of this polynomial at the cell's own mean state. This is the Jacobian entry $\partial(dx_j/dt)/\partial x_k$ evaluated at the mean of the cell's observed protein levels. An arrow is green when this entry is positive and purple when it is negative, and its width grows with the entry's magnitude. When the derivative changes sign over the range of protein $k$ that the cell actually visited, so that the regulator activates at some levels and inhibits at others, the arrow is dashed, and its color reports the sign at the mean state only.

#### C.3 Multidimensional scaling to visualize network diversity

The embedding of Figure 7B places all 207 cells in a plane so that cells with similar estimated dynamics are close. Every PRNODE fit yields two fitted systems, one per cell of the pair, and a cell that belongs to several pairs, a mother of several daughters or a daughter who later becomes a mother, has several fitted systems. We compare any two fitted systems, whether or not they come from the same pair, by the flow-map similarity (6) used for inheritance: each system is linearized at its own cell's mean observed state, as in the previous subsection, after every protein is put on one population scale (its standard deviation over all timepoints of all 207 cells) so that systems fitted to different pairs are comparable, and the dissimilarity between systems $a$

and $b$ is $d(a,b) = \|\Phi_a - \Phi_b\|_F / (\|\Phi_a - I\|_F + \|\Phi_b - I\|_F)$, one minus the similarity, where $\Phi_a = \exp(hA_a)$ is the one-timepoint flow map of system $a$ at $h = 0.2$ hours. This is computed for every pair of fitted systems by stacking the vectorized $\Phi_a - I$ as rows, taking their Euclidean distances with `stats::dist()`, and dividing by the sum of the two row norms, which gives a matrix with entries in $[0, 1]$ and zeros on the diagonal. This yields a 360-by-360 matrix over the fitted systems, two from each of the 180 pairs.

This matrix is then collapsed to the 207 cells by averaging $d(a, b)$ over all pairs of systems $a$ of one cell and $b$ of the other, with the diagonal set to zero, and embedded in two dimensions by classical multidimensional scaling, `stats::cmdscale()` applied to `stats::as.dist()` of the cell-by-cell matrix with `k = 2` and `eig = TRUE`.

##### C.4 Global network estimation across all yeast cells simultaneously

Figure 7C compares the per-pair PRNODE fits with one network fitted to all $N = 207$ cells at once, the multiple-trajectory version of GRADE [8], in which every cell is a realization of the same ODE and differs from the others only in its initial condition. For protein $j$ the global estimate solves

$$\min_{\theta_j \in \mathbb{R}^{pq}} \frac{\omega}{2} \sum_{c=1}^{N} \left\| \widetilde{y}_j^{(c)} - \sum_{k=1}^{p} \widetilde{\Psi}_k^{(c)} \theta_{jk} \right\|_2^2 + \lambda\sqrt{q} \sum_{k=1}^{p} \left\| \omega^{1/2} \left[ \widetilde{\Psi}_k^{(1)\top}, \ldots, \widetilde{\Psi}_k^{(N)\top} \right]^\top \theta_{jk} \right\|_2, \tag{S6}$$

where $\widetilde{y}_j^{(c)}$ and $\widetilde{\Psi}_k^{(c)}$ are cell $c$'s profiled response and integrated basis of Section 3.1, now with only the initial condition profiled out of each cell and one constant term shared by all cells, and $\omega$ is one common weight, the timepoint-weighted pool of the cells' inverse noise variances. This is the scaffold block of PRNODE's stacked design (S2) extended from two cells to $N$, with the deviations removed, and a check on one pair confirms that the code reproduces PRNODE with the deviations switched off. Everything else matches the per-pair pipeline: the quadratic basis $q = 2$, every protein scaled by its population standard deviation, each cell smoothed at the same smoothing level, the same group-lasso solver, and the smoothing level and $\lambda$ chosen together by the known-variance BIC of Appendix B (the BIC chooses the roughest smoothing level and a network of 26 off-diagonal edges).

The error of a fit for cell $c$ is the mean over its timepoints and the six proteins of the squared difference between the fitted trajectory, solved forward from the cell's estimated initial condition, and the observed values, in units of each protein's population variance; for the per-pair fits, a cell that belongs to several pairs is scored by the mean of its errors across those fits.

##### C.5 Pairing daughters with unrelated mothers

Figure 8B asks whether a daughter's true mother is the mother whose network is most similar to hers. We take the 67 first-generation daughters, the daughters of the 26 founder cells, with more than 12 timepoints, and pair each one with each of the 25 founders of the other families, a founder being the one cell of its family that is present from the first timepoint. Every unrelated pairing is fitted exactly as a true pair, from the raw timepoints with the same basis $q = 2$, the same BIC over the smoothing level and $\lambda$, and the same inheritance estimate $\widehat{\tau}$, for 1,675 fits in all. For daughter $D_i$ with true mother $M_i$ and unrelated founders $m_1, \ldots, m_{25}$, the rank of the true mother is

$$\text{rank}_i = 1 + \#\big\{ m \in \{m_1, \ldots, m_{25}\} : \widehat{\tau}(m, D_i) > \widehat{\tau}(M_i, D_i) \big\}, \tag{S7}$$

so that rank 1 means no founder reads as more similar to the daughter than her own mother, rank 26 means every founder does, and smaller ranks show more inheritance. Under no inheritance the pairings are exchangeable and the rank is uniform on $\{1, \ldots, 26\}$, with mean 13.5, and $\text{rank}_i/26$ is the exact permutation $p$-value of daughter $i$, whose smallest attainable value is $1/26 \approx 0.04$. Figure 8B shows the histogram of the 67 ranks against this uniform reference. The 1,675 unrelated-mother estimates are also the null band of Figure 8A: its dashed line is their median and its band runs from their 2.5th to their 97.5th percentile.

#### D Pseudocode of PRNODE

Algorithm S1 summarizes the procedure for one mother-daughter pair, with $n_c$ the number of timepoints of cell $c$ ($n_M$ and $n_D$ in the main text).

**Algorithm S1:** PRNODE for one mother-daughter pair.

**Input :** Observed trajectories $\{(t_i^{(c)}, y_i^{(c)})\}_{i=1}^{n_c}$ for $c \in \{M, D\}$ (with $n_M$ and $n_D$ timepoints); polynomial degree $q$; multipliers $\mu_M = \mu_D = 1$; smoothing grid $\{0, 0.25, 0.5, 0.75\}$; size of $\lambda$ candidate grid (default $n_\lambda = 80$)

**Output:** Networks $\widehat{A}^{(S)}, \widehat{A}^{(M)}, \widehat{A}^{(D)}$, the coefficient estimates, the selected smoothing parameter and $\lambda$, and the inheritance $\widehat{\tau}$.

1 Divide each protein by its pooled standard deviation $s_j$ over both cells, and compute the common loss weight $\omega$ from the second-difference noise estimates;

2 **for** *each smoothing parameter* `spar` *in the grid* **do**

3 Smooth every protein's series in each cell with a smoothing spline at `spar` and compute the integrated basis $\widehat{\Psi}_k^{(c)}(t_i^{(c)})$ by the trapezoidal rule;

4 Profile out each cell's initial condition and constant term, and stack the two cells into the weighted design (S2) with $3p$ column groups;

5 Compute $\lambda_{\max}$ and the $n_\lambda$ log-spaced values of $\lambda$ from $\lambda_{\max}$ to $10^{-4}\lambda_{\max}$;

6 **for** *each protein* $j$ **do**

7 Fit the group-lasso path of (S3) over the $\lambda$ values by cyclic block coordinate descent with warm starts;

8 **end**

9 For each $\lambda$, refit the coefficients for the selected groups of every protein by least squares and evaluate BIC($\texttt{spar}, \lambda$) of (S4);

10 **end**

11 Select $(\widehat{\texttt{spar}}, \widehat{\lambda})$ as the combination of $\{\texttt{spar}, \lambda\}$ that minimizes BIC($\texttt{spar}, \lambda$);

12 Record the $\{\widehat{\theta}_{jk}^{(S)}, \widehat{\delta}_{jk}^{(M)}, \widehat{\delta}_{jk}^{(D)}\}_{j,k=1}^{p}$ that minimized (S3) for the selected $(\widehat{\texttt{spar}}, \widehat{\lambda})$;

13 Undo the protein scaling, set $\widehat{\theta}_{jk}^{(c)} = \widehat{\theta}_{jk}^{(S)} + \widehat{\delta}_{jk}^{(c)}$ for $c \in \{M, D\}$, and record $\widehat{A}_{jk}^{(c)} = 1\{\widehat{\theta}_{jk}^{(c)} \neq 0\}$ for $c \in \{S, M, D\}$;

14 Compute $\widehat{\tau}$ as the plug-in of (6) at the estimated coefficient matrices. (See Appendix B.4 for additional details if $q > 1$);

15 **return** $\widehat{A}^{(S)}, \widehat{A}^{(M)}, \widehat{A}^{(D)}, \widehat{\tau}$

#### E Theoretical properties of PRNODE

##### E.1 Mathematical theorems related to the tuning of PRNODE

With $\widetilde{\mathbf{y}}_j$ and the column groups $X_{S,k}$, $X_{M,k}$, and $X_{D,k}$ of the stacked design (S2), write $\|v\|_{g,k} = \|X_{g,k}v\|_2$ for $g \in \{S, M, D\}$, the norm of the fitted contribution of one group in (S3), and $\mu_S = 1$.

**Proposition 1 (Empty-network penalty).** *Assume that each $X_{g,k}$ has full column rank. The all-zero vector minimizes* (S3) *for protein $j$ if and only if*

$$\lambda \;\geq\; \lambda_{\max,j} = \max_{k\in\{1,\ldots,p\}} \max_{g\in\{S,M,D\}} \frac{\big\|(X_{g,k}^\top X_{g,k})^{-1/2} X_{g,k}^\top \widetilde{\mathbf{y}}_j\big\|_2}{\sqrt{q}\,\mu_g}, \tag{S8}$$

*and it is the unique minimizer when the inequality is strict.*

The numerator in (S8) is the norm of the least-squares projection of $\widetilde{\mathbf{y}}_j$ onto the columns of one group, so $\lambda_{\max,j}$ is the largest fitted contribution that a single regulator can make on its own in any of the three components, and the tuning path starts at $\lambda_{\max} = \max_j \lambda_{\max,j}$, where all three networks are empty.

**Proposition 2 (Sharing between the cells).** *If $\mu_M^2 + \mu_D^2 < 1$, then every minimizer of* (S3) *has $\widehat{\theta}_{jk}^{(S)} = 0$ for all $k$, and $\widehat{\theta}_j^{(c)} = \widehat{\delta}_j^{(c)}$ minimizes the cell-c terms of* (S3) *on their own, so that the estimator decouples into one group-lasso fit per cell and neither cell's data enter the other's estimate.*

A common multiplier $\mu_M = \mu_D = \mu < 1/\sqrt{2}$ therefore shares nothing, whatever the data. At $q = 1$ the opposite holds at $\mu_M = \mu_D = 1$: whenever the two cells' estimated coefficients of a regulator have the same sign, the scaffold takes the one of smaller magnitude and that cell's deviation is zero, whatever the two cells' lengths and noise levels, while two coefficients of opposite sign leave the scaffold empty:

**Proposition 3 (Sharing rule at $q = 1$).** *Let $q = 1$, and write $a_{c,k} = \|\widetilde{\Psi}_k^{(c)}\|_2$ for $c \in \{M, D\}$ and $a_{S,k} = (a_{M,k}^2 + a_{D,k}^2)^{1/2}$. Suppose that $\widehat{\theta}_{jk}^{(M)}$ and $\widehat{\theta}_{jk}^{(D)}$ are nonzero with the same sign. If $a_{S,k} < \mu_M a_{M,k} + \mu_D a_{D,k}$, then $\widehat{\theta}_{jk}^{(S)} \neq 0$ and equals one of the two cells' coefficients, so that cell's deviation is zero; if $a_{S,k} > \mu_M a_{M,k} + \mu_D a_{D,k}$, then $\widehat{\theta}_{jk}^{(S)} = 0$. At $\mu_M = \mu_D = 1$ the scaffold equals the coefficient of smaller magnitude, and two coefficients of opposite sign give $\widehat{\theta}_{jk}^{(S)} = 0$.*

In other words, because $(a_{M,k} + a_{D,k})/\sqrt{2} \leq a_{S,k} < a_{M,k} + a_{D,k}$ whenever both cells have a nonzero column for regulator $k$, a common multiplier $\mu_M = \mu_D = \mu \leq 1/\sqrt{2}$ never places an edge in the scaffold, in agreement with Proposition 2, while $\mu = 1$ places every edge that the two cells estimate with the same sign in the scaffold, whatever the two cells' lengths and noise levels. Due to this proposition, we therefore hold $\mu_M = \mu_D = 1$ by default, so that every pair shares its common edges through the scaffold on the same terms and the inheritance measure of Section 3.2 stays comparable across pairs. The smoothing parameter and $\lambda$ are then chosen from the data by the BIC of Appendix B.

##### E.2 The null value of the inheritance measure

Two unrelated networks do not have an inheritance of zero. Write $\tau_h(A^{(M)}, A^{(D)})$ for the measure (6) at sampling interval $h$ and coefficient matrices $A^{(M)}, A^{(D)} \in \mathbb{R}^{p\times p}$, the matrices $J^{(c)}$ of Section 3.2 at $q = 1$, and

$$\tau_0(A^{(M)}, A^{(D)}) = 1 - \frac{\|A^{(M)} - A^{(D)}\|_F}{\|A^{(M)}\|_F + \|A^{(D)}\|_F} \tag{S9}$$

for its first-order version. The following proposition says that the measure is a normalized distance between the two generators of the dynamics at small $h$ and that two networks of equal size with nothing in common sit at $1 - 1/\sqrt{2}$.

**Proposition 4 (Null value of the inheritance).** *Let $q = 1$ and $A^{(M)}, A^{(D)} \in \mathbb{R}^{p\times p}$ with $\|A^{(M)}\|_F + \|A^{(D)}\|_F > 0$.*

*(a) As $h \to 0$, $\tau_h(A^{(M)}, A^{(D)}) = \tau_0(A^{(M)}, A^{(D)}) + O(h)$.*
*(b) If $\|A^{(M)}\|_F = \|A^{(D)}\|_F$ and $\mathrm{tr}(A^{(M)\top}A^{(D)}) = 0$, then $\tau_0(A^{(M)}, A^{(D)}) = 1 - 1/\sqrt{2} \approx 0.293$.*

Part (b) says that two networks of the same total strength whose coefficient vectors are orthogonal are as different as two networks of that size can be without pointing in opposite directions, and the measure places them at 0.29 rather than at 0. A common structure that both cells carry, such as the self-decay of every protein, raises the null above 0.29: in our simulated populations the null sits near 0.33, and the unrelated-mother null of Section 5 has a median of 0.26, below 0.29 because the two fitted systems of an unrelated pair differ in size as well as in direction. This is why a pair's estimate is read against the empirical null built from unrelated pairs in Section 5, rather than against zero. At $q > 1$ the same statements hold with $A^{(c)}$ replaced by the Jacobian of cell $c$'s fitted field at its own mean state, which is what (6) exponentiates. The proof is in Appendix H.

### F Additional simulation results

#### F.1 Design of the simulated data

Here we provide further details on how we simulate networks and noisy time series data for the mother-daughter pairs in our simulation study. For each distinct mother replicate, we start by choosing a self-degradation rate for each protein uniformly on $[-2.5, -0.5]$, which is the coefficient placed on the linear term for that protein's impact on its own derivative in its ODE system. For the mothers in the cubic regime, the quadratic and cubic coefficients of the protein for its own derivative are the self-degradation rate divided by 2! and 3!, respectively. Then, we draw 6 off-diagonal edges of the network that will belong to the scaffold and 2 that are meant to be the mother's deviation in antisymmetric pairs. For the monomial regimes, the magnitude of the edge coefficients $m$ is chosen uniformly on $[4\pi, 10\pi]$ and the ODE system has coefficients $+m$ and $-m$ at positions $(j, k)$ and $(k, j)$ for the chosen off-diagonal edges. This ensures that each pair is a damped rotation that completes two to five cycles on the $[0, 1]$ time grid and the integrated designs stay well conditioned. For the mother networks in the cubic regime, the coefficients for the quadratic and cubic coefficients are also drawn uniformly on $[4\pi, 10\pi]$ and then divided by the factorial of their order so they scale appropriately. After determining the mother network, we proceed to drawing the daughter networks. For the high inheritance pairs, the daughter is first assigned the exact same ODE system as her mother (scaffold and mother's two deviations), but then half of the shared edges are made unequal by scaling the daughter's coefficients by a factor chosen uniformly on $[0.5, 2]$. At the other three levels, the daughter keeps only the 6 designated scaffold edges and then a pair of antisymmetric edges is drawn so that she has 2 edges that deviate from her mother. Again, half of the daughter's shared edges are made unequal through scaling them by a factor chosen uniformly on $[0.5, 2]$. Then, the daughter's deviation from the scaffold is multiplied by 1, 2, or 4 (moderate, mild, and low), which lowers the inheritance (6) because the strength of the daughter's differing edges grows relative to strength of the edges shared with its mother. Consequently, there is some overlap in the ranges of true inheritance values found among pairs of each inheritance level because the multiplier can only indirectly impact the true inheritance value via the size of the daughter's deviation.

We solve each pair's system numerically from a standard normal initial state to obtain the mother's noiseless trajectory values at the 48 equally spaced timepoints on $[0, 1]$, and the daughter's noiseless observations are set as the mother's final $n_D$ timepoints. We then draw Gaussian noise terms as $\epsilon \sim N(0, Ls_j r_j)$, where $L$ is a noise level, $s_j$ is the standard deviation of the mother's 48 noiseless observations, and $r_j$ is a scaling factor for protein $j$ so that each protein has slightly different noise distributions. The noise level is set as $L = 0.1$ for the low noise and misspecified regimes and $L = 1.0$ at high noise, and $r_j$ is drawn log-uniformly between 1/1.5 and 1.5.

#### F.2 Additional simulation results

Here we provide additional results from our simulation studies that were not presented in the main text.

First, we present results for the MCC of the mother network estimates from all four methods for high and low inheritance pairs across all regimes (Figure S2). The three methods that fit each cell on its own give the same mother estimates at the two inheritance levels as the mother does not change across daughter lengths, so only PRNODE's results differ across inheritance levels due to its joint estimation approach. PRNODE outperforms the other methods in the low and high noise regime at every length. However, the kernel ODE recovers the mother's network better than PRNODE in the misspecified regime. This tracks with our results for the daughter network estimates as well. The kernel ODE's nonparametric basis likely makes its estimates more robust to the misspecified cubic field.

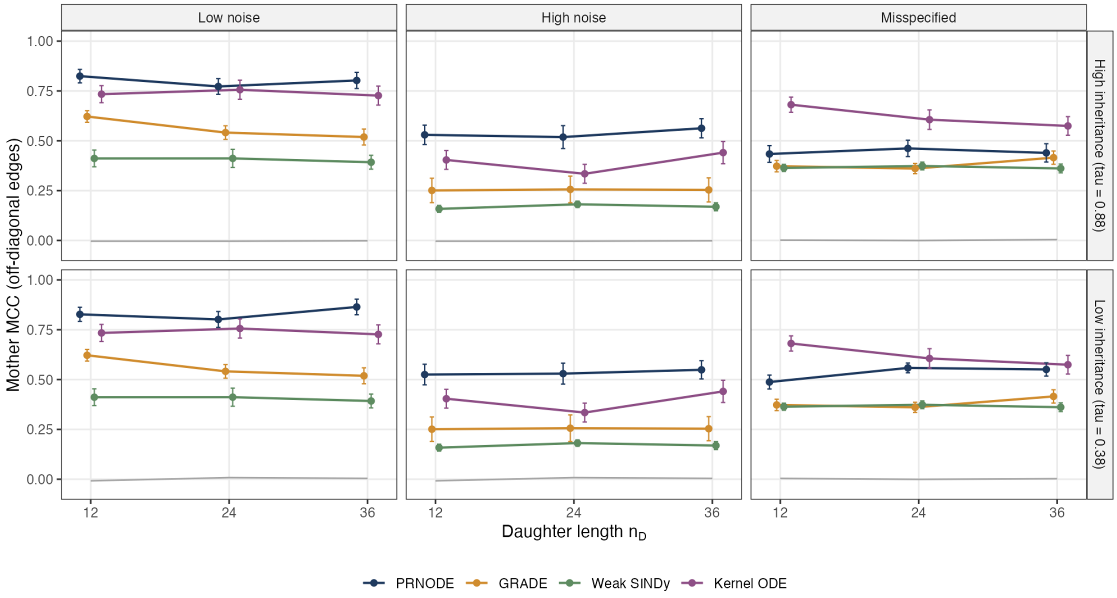


**Figure S2.** Mother's network recovery (MCC over the 30 off-diagonal edges) results for low and high inheritance. MCC is given on the y-axis, daughter's length on the x-axis, and the methods are indicated by color. The inheritance levels are across the rows, regimes across columns, and the points are means over 20 replicates with ±1 standard error. The gray line is the chance level.

We also provide the network recovery results for all methods for the mild and moderate inheritance pairs across all regimes (Figure S3). The results here track those for the other inheritance levels, where PRNODE outperforms all other methods for the low and high noise regime and outperforms all methods aside from the kernel ODE in the misspecified regime.

We also provide the comparison of inheritance estimates produced by PRNODE's network fits and GRADE's network fits for all regimes when the daughters have more observations (Figure S4). PRNODE's inheritance estimates are more strongly correlated with the truth than GRADE's across every regime at the longer daughter lengths as well. GRADE's inheritance estimates are still mostly 0 or 1 in the misspecified regime even with more daughter observations. In contrast, PRNODE's inheritance estimates improve as the number of daughter observations grows.

Our tuning procedure investigation in the main text is augmented by comparison of the MCC for networks picked by our tuning procedure compared to the best possible networks produced across the full tuning parameter grid (Figure S5). We designate the oracle network estimate as the one produced by choosing the $\lambda$ and smoothing parameter combination along the joint grid for both that has the highest MCC for its estimated networks. The oracle $\lambda$ is selected based on knowledge of the true network, so it is an upper bound on what any tuning rule applied to PRNODE's path could achieve. PRNODE's estimated networks only narrowly underperform the oracle estimate for both daughter and mother network recovery across the low and high noise regimes. The gap between PRNODE and the oracle widens in the misspecified regime.

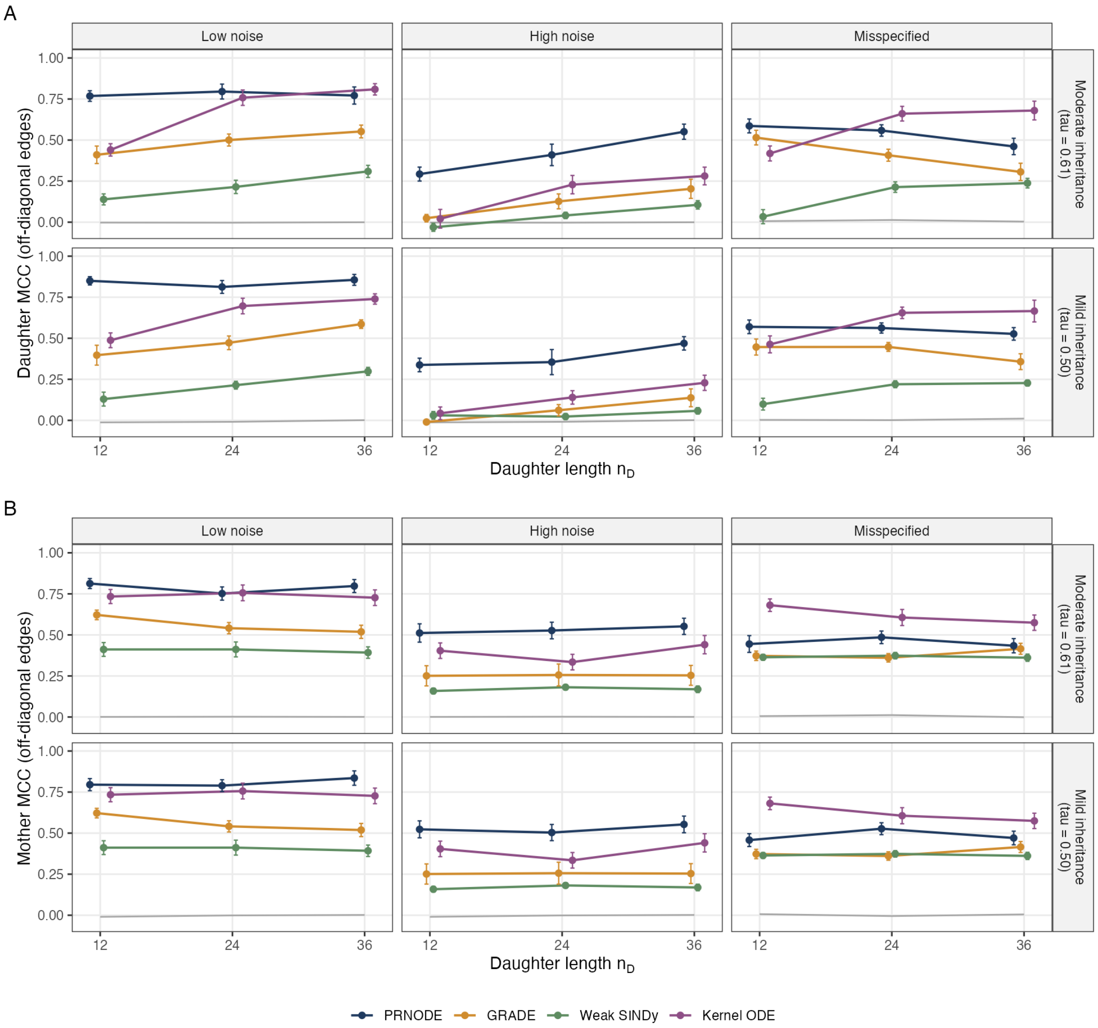


**Figure S3.** Network recovery (MCC over the 30 off-diagonal edges) results for both daughter (A) and mother (B) for mild and moderate inheritance pairs. MCC is given on the y-axis, daughter's length on the x-axis, and the methods are indicated by color. The inheritance levels are across the rows, regimes across columns, and the points are means over 20 replicates with ±1 standard error. The gray line is the chance level.

We investigate the power of our permutation test for inheritance as well (Figure S6). For this, we perform the inheritance test for all high inheritance mother-daughter pairs in the low noise regime of our simulation where $n_D = 12$. For each daughter, the test compares the inheritance estimated with her true mother with the inheritance estimated with 19 unrelated mothers, so that the smallest possible p-value is $1/20 = 0.05$. The true mother ranks first for 18 of the 20 daughters, so the test rejects at level 0.05 for 18 daughters and at level 0.1 for 19 of them. The remaining daughter for which the test does not reject has a p-value of 0.45. This result suggests that even with few daughter observations, our permutation test is well-powered for a correctly specified model and low noise observations.

We also provide a comparison of the run times for PRNODE and all the methods we compared in our simulation study. PRNODE's median time is 46 seconds, below that of weak SINDy and the kernel ODE and above that of GRADE, and its mean is 194 seconds (Table S1). PRNODE's runtime falls as the daughter's series grows, from a mean of 464 seconds at $n_D = 12$ to 27 seconds at $n_D = 36$, which is the opposite of what the size of the problem suggests. This is likely a consequence of the cost of the search over the smoothing level and $\lambda$.

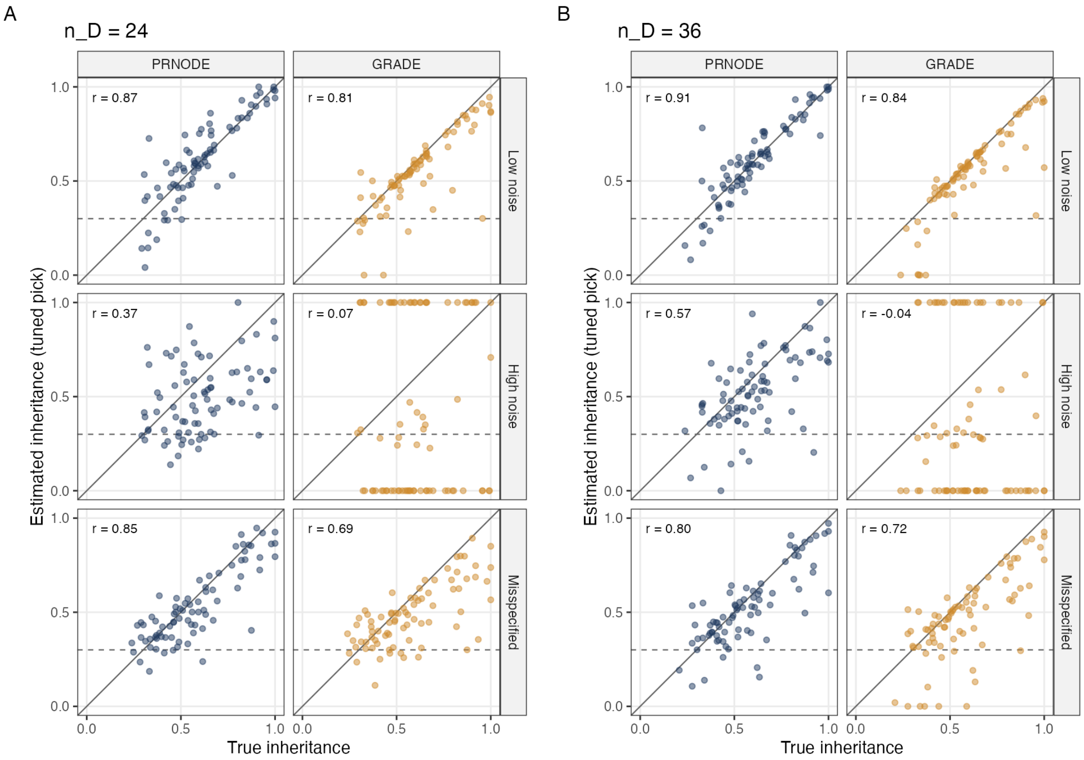


**Figure S4.** Estimated versus true inheritance for daughters with $n_D = 24$ (A) and $n_D = 36$ (B) across PRNODE and GRADE. The true inheritance is given on the x-axis and estimated inheritance on the y-axis. The methods are faceted across the columns and regimes across the rows. The solid line is the identity and the dashed line is the inheritance of unrelated networks, about 0.3. The correlation is printed in each panel.

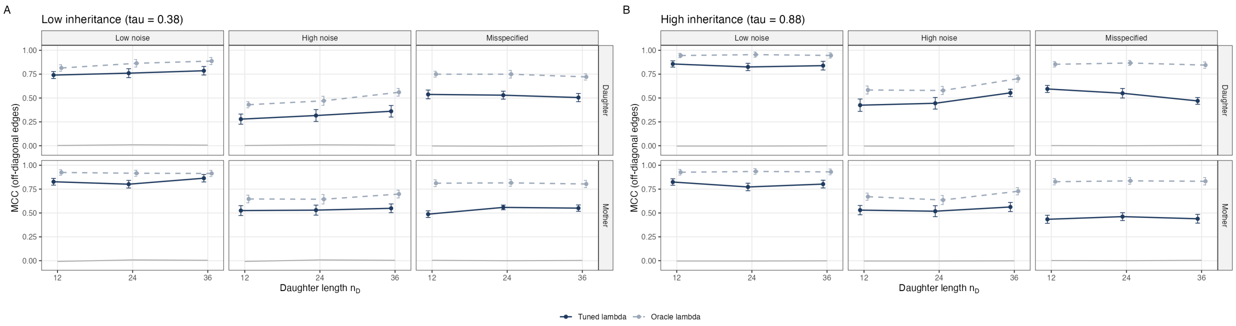


**Figure S5.** MCC of PRNODE at its tuned $\lambda$ (solid) and at the oracle $\lambda$ (dashed), for the daughter and mother at low (A) and high (B) inheritance. Points are means over 20 replicates with $\pm 1$ standard error, and the gray line is the chance level.

### G Additional details of yeast data

#### G.1 Details about experimental setup and image preprocessing

The live-cell imaging data was collected using the conditions described in Arguello et. al. [2], in which diploid budding yeast cells growing in a microfluidic chamber under a constant nutrient supply are imaged every 12 minutes for 48 timepoints (9.4 hours), in a phase contrast channel and six fluorescence channels, one per tagged protein. The six proteins are chosen because their nuclear levels together read a cell's position in the division cycle and its response to the environment, and the six fluorescent tags are chosen for their spectral separation, which is what limits the number of proteins that can be followed at once. Three report

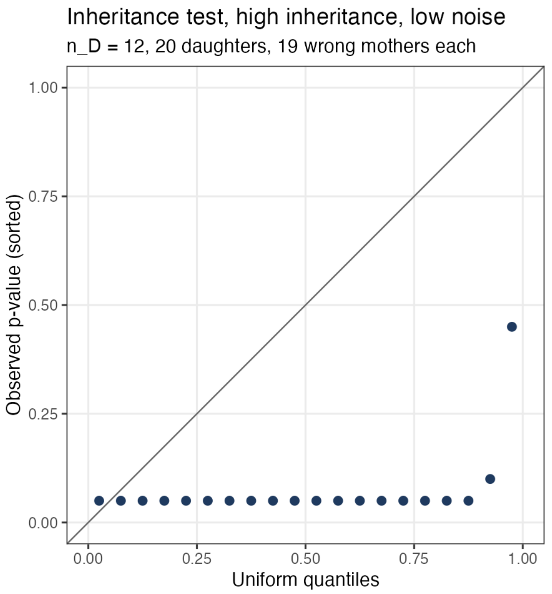


**Figure S6.** Quantile-quantile plot of the permutation test's p-values against uniform quantiles, for the 20 daughters at the highest inheritance level, low noise, and $n_D = 12$.

**Table S1.** Average runtime in seconds per dataset, mean (median) over the 240 datasets at each daughter length $n_D$ and over all 720. Times come from a cluster with one CPU per task. Weak SINDy's time is the time of the call over its cells, with 5 subdomain seeds. GRADE's extra refit for the inheritance estimate is not included.

| Method | $n_D = 12$ | $n_D = 24$ | $n_D = 36$ | Overall |
|---|---|---|---|---|
| PRNODE | 464 (276) | 93 (37) | 27 (14) | 194 (46) |
| GRADE | 93 (43) | 42 (22) | 46 (20) | 60 (25) |
| Weak SINDy | 204 (212) | 203 (213) | 203 (212) | 204 (212) |
| Kernel ODE | 544 (551) | 566 (580) | 571 (584) | 560 (571) |

the division cycle: Whi5 (tagged with mKO$\kappa$), the transcriptional repressor of the G1/S regulon, which is nuclear in G1 and leaves the nucleus at the commitment to division; Clb2 (mNeonGreen), the main mitotic B-type cyclin, which accumulates in the nucleus before anaphase and is degraded at mitotic exit; and Cdc10 (mCyOFP1), one of the septins that form the ring at the mother-bud neck after commitment. Three report the response to nutrients and stress: Xbp1 (mRuby3), a stress-induced repressor that accumulates in the nucleus on starvation and promotes quiescence [15,2]; Stb3 (mTFP1), a repressor of ribosome biogenesis genes that enters the nucleus when nutrients fall [14]; and Tup1 (mNeptune2.5), the general corepressor recruited to glucose-repressed and stress genes [26]. The protein values are summary statistics of each cell's segmented image at each timepoint: for the five nuclear proteins, the fluorescence concentration in the nucleus, and for Cdc10, the standard deviation of the fluorescence at the cell periphery, which is high when the septin ring is assembled and low otherwise, so that all six values rise and fall with the cell's state rather than with its size.

Cells are segmented and tracked by FIEST [11], a convolutional-network pipeline that delineates each cell at every timepoint, links the masks across timepoints into one track per cell, and records each bud's birth timepoint and its mother, which defines the lineages. Of the 239 tracked cells, we keep the 207 whose tracks can be assigned to one of the 26 founder cells present at the first timepoint and whose mother is identified, which gives 180 mother-daughter pairs; 99 distinct mothers appear, since a mother contributes one pair per daughter, and the mothers are founders in 74 pairs, first-generation daughters in 72, and later generations in 34. A daughter's series starts at her birth timepoint and runs to the end of the movie, so its length depends on when she was born: the median daughter has 21 timepoints, the shortest 5, and 50 of the 180 daughters have 12 or fewer. A timepoint at which a cell's mask is missing is filled by linear interpolation within the cell's own series, and time is measured in hours from each cell's birth, so that a mother and her daughter are fitted on their own clocks.

#### G.2 Details about analyses performed

Several details of the tests behind Figures 7 and 8 are left out of the main text.

- **Figure 7C (global fit).** The $p$-value is a paired Wilcoxon signed-rank test of the 207 per-cell errors under the global model against those under the per-pair fits. A cell in several pairs is scored by the mean of its per-pair errors.
- **Figure 7D (Jaccard).** Each network is the set of nonzero off-diagonal edges, and the Jaccard similarity is the size of the intersection over the size of the union of the two sets. The "same mother" group pairs the two networks estimated for one mother in two different pairs (83 comparisons from the mothers with at least two daughters), and the "different mother" group pairs networks of different mothers (2,618); the "true pair" group compares a daughter's network with her own mother's (74 daughters who are also mothers, so that each is estimated in a pair as a daughter and in a pair as a mother), and the "fake pair" group compares each such daughter with every other mother (5,236). The $p$-values are two-sided Wilcoxon rank-sum tests. The comparisons within a group are not independent, since one cell enters many of them, so the $p$-values are descriptive.
- **Figure 8A (null band).** The band is the 2.5th to the 97.5th percentile, and the dashed line the median, of the 1,675 estimates from first-generation daughters paired with an unrelated founding mother of Appendix C. Of the 180 true pairs, 16 lie above and 11 below, against 4.5 expected on each side.
- **Figure 8B (rank of the true mother).** The dashed reference is the mean rank, 13.5, under a uniform rank; the summary tests are a one-sided binomial test of 42 of 67 daughters in the upper half ($p = 0.02$) and a Wilcoxon signed-rank test of the 67 percentiles against one half ($p = 0.03$). Daughters with 12 or fewer timepoints are excluded because their fits are too unstable for 25 refits to be informative.
- **Figure 8C (what tracks inheritance).** The left panel's association is a Spearman correlation between the estimate and the number of shared edges whose Jacobian entries at the two cells' mean states agree in sign, over the 180 pairs, with the Spearman $p$-value; the count tracks the estimate partly because longer daughters have both more edges and higher estimates (Spearman 0.24 between the estimate and the daughter's length). In the right panel the four states of one edge are read from the sign of the Jacobian entry at each cell's mean state, the $p$-value is a Kruskal-Wallis test across the four states, and 30 off-diagonal edges are examined, of which Whi5 to Cdc10 has the smallest $p$-value.

#### G.3 Additional results about yeast cells

#### G.4 The imaged regulators in the division cycle

The nuclear levels of a few cell-cycle regulators are enough to read a cell's position in the cycle and its entry into proliferation (cell division) or quiescence from a single image [11,2]. Among the proteins we image, Whi5 and Clb2 act in opposite phases of the cycle: nuclear Whi5 represses the transcription of the G1/S genes and so holds the cell in G1 until Cdk1 activity drives it from the nucleus [25], after which the mitotic cyclin Clb2 accumulates and its Clb2-Cdk1 complex drives the cell through mitosis, and Clb2's degradation at mitotic exit is what lets Whi5 return to the nucleus [1,7,11]. A third protein ties into this loop from outside the cycle: the stress-induced repressor Xbp1 promotes the nuclear accumulation of Whi5 and thereby entry into quiescence [2], so that an environmental signal changes the Whi5 side of the loop.

#### G.5 Biology behind the most frequent shared edges

The edges most often shared by a mother and her daughter (Figure 7E) are, almost without exception, edges among the three cell-cycle proteins and from them to the stress repressors, and the direction PRNODE assigns to each at the cells' mean states agrees with what is known of the division cycle.

– *Whi5 inhibits Clb2 and Clb2 activates Whi5.* Nuclear Whi5 represses the G1/S transcription factor SBF, and through it the G1/S cyclins whose Cdk1 activity starts the cascade that ends in Clb2 accumulation [25]; Cln-Cdk1 phosphorylation drives Whi5 out of the nucleus at Start [25], after which Clb2-Cdk1 keeps SBF off through mitosis [1], and Clb2's degradation by the anaphase-promoting complex at mitotic exit lets Whi5 re-enter the nucleus [7]. The fitted edge from Clb2 to Whi5 is activating in 81% of the cells that are estimated to have this edge, which reads the return of Whi5 as Clb2 falls, and the edge from Whi5 to Clb2 is inhibiting in 72%, which reads the hold on the cycle while Whi5 is nuclear.
– *Whi5 inhibits Cdc10.* The septin ring at the incipient bud site assembles only after Start once Whi5 has left the nucleus, and is disassembled at cytokinesis [16]. Therefore, the Cdc10 periphery signal rises when nuclear Whi5 is low and falls as Whi5 returns; the edge is inhibiting in 81% of the cells that are estimated to have this edge.
– *Clb2 activates Xbp1.* Xbp1 can be transcribed in G2/M with a peak at the time of Clb2 accumulation [2] and is also induced by stress [15]; the fitted edge is activating in 80% of the cells that are estimated to have this edge.
– *Clb2 and Cdc10.* An edge from Clb2 to Cdc10 is less frequent (27%). Clb2-Cdk1 is recruited to the bud neck in anaphase through a docking motif in Bud3 [19], and the septin-associated protein Syp1, which binds Cdc10, accelerates ring assembly and disassembly [20], so the two proteins meet at the bud neck. No Clb2-dependent modification of Cdc10 has been shown directly as far as we know, so we interpret this edge as the two proteins reporting coupled stages of the same mitotic program rather than as a direct molecular interaction.

Figure S7 overlays the mother-daughter pairs on the embedding of Figure 7B. Every cell is a point, founders as squares and daughters as circles, and each pair is a segment from the mother to the daughter, gray for most pairs and colored for the 18 pairs nearest an inheritance estimate of 0, the 18 nearest 0.3 (the null value of Proposition 4), and the 18 nearest 1 (the 18 highest, from 0.57 upward). The segments of the low-inheritance pairs cross the plane, often from one side to the other, whereas the segments of the high-inheritance pairs are short, and the pairs near 0.3 sit in between; over all pairs the distance in the plane and the inheritance estimate have a Spearman correlation of $-0.42$.

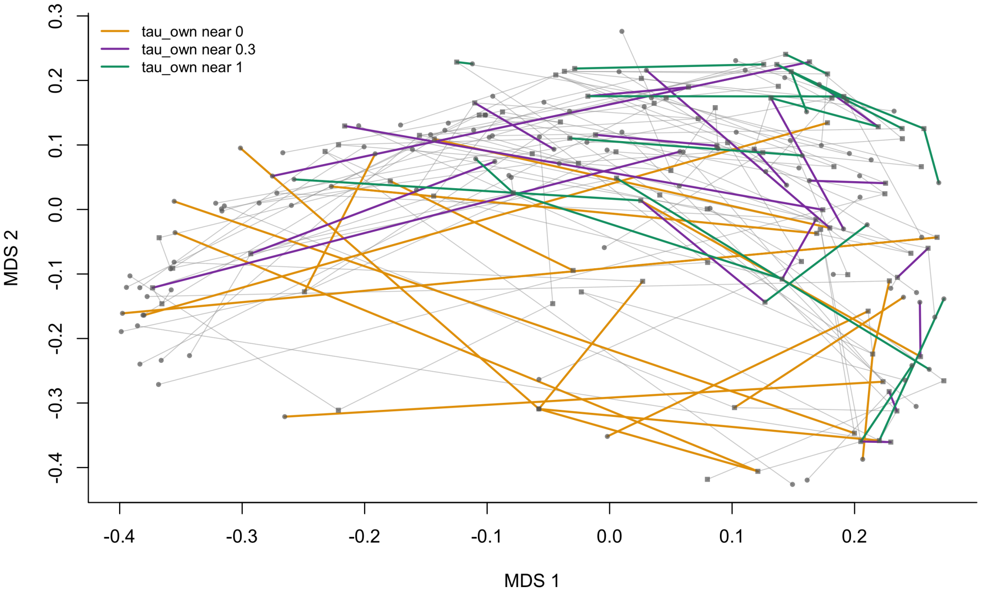


**Figure S7. Mother-daughter pairs on the embedding of the 207 cells.** Each point is one cell at its position in the multidimensional scaling of Figure 7B (squares: founders; circles: daughters), and each segment joins a mother to her daughter. The 18 pairs with the lowest inheritance estimates ("near 0"), the 18 nearest the null value of 0.3, and the 18 highest ("near 1") are colored; the remaining pairs are gray. Pairs with lower inheritance are placed further apart in the plane.

Figure S8 extends the right panel of Figure 8C from one edge to all 30 off-diagonal edges. For each edge, the 180 pairs are split into four states by the signs of the edge's Jacobian entries at the two cells' mean states, present in both cells with the same sign, present in exactly one cell, present in neither, or present in both with opposite signs, and the mean inheritance estimate within each state is plotted, with the edges ordered from top to bottom by how often they are shared. The pattern of Whi5 to Cdc10 recurs across the frequent edges: for the 13 edges shared by at least 28% of pairs, the pairs with the same sign have the highest mean, between 0.33 and 0.40, the pairs in which neither cell has the edge sit near or below the overall mean of 0.30, and the pairs with opposite signs, which are few, sit lowest. For the rare edges at the bottom of the figure the four means converge toward 0.30, because an edge that few pairs share cannot separate them. Two caveats keep this suggestive rather than definitive. First, a shared same-sign edge raises the inheritance estimate whether the edge is specific to that lineage or common to every yeast cell, so the figure does not separate what is inherited from what is generic, and the most frequent edges are, by construction, those every cell tends to have. Second, the means of the same-sign groups, 0.33 to 0.40, lie inside the null band of Figure 8A, whose 97.5th percentile is 0.57, so a typical pair in the same-sign state is not individually distinguishable from an unrelated pairing; what the figure shows is a consistent shift of group means, not a set of pairs that clear the null.

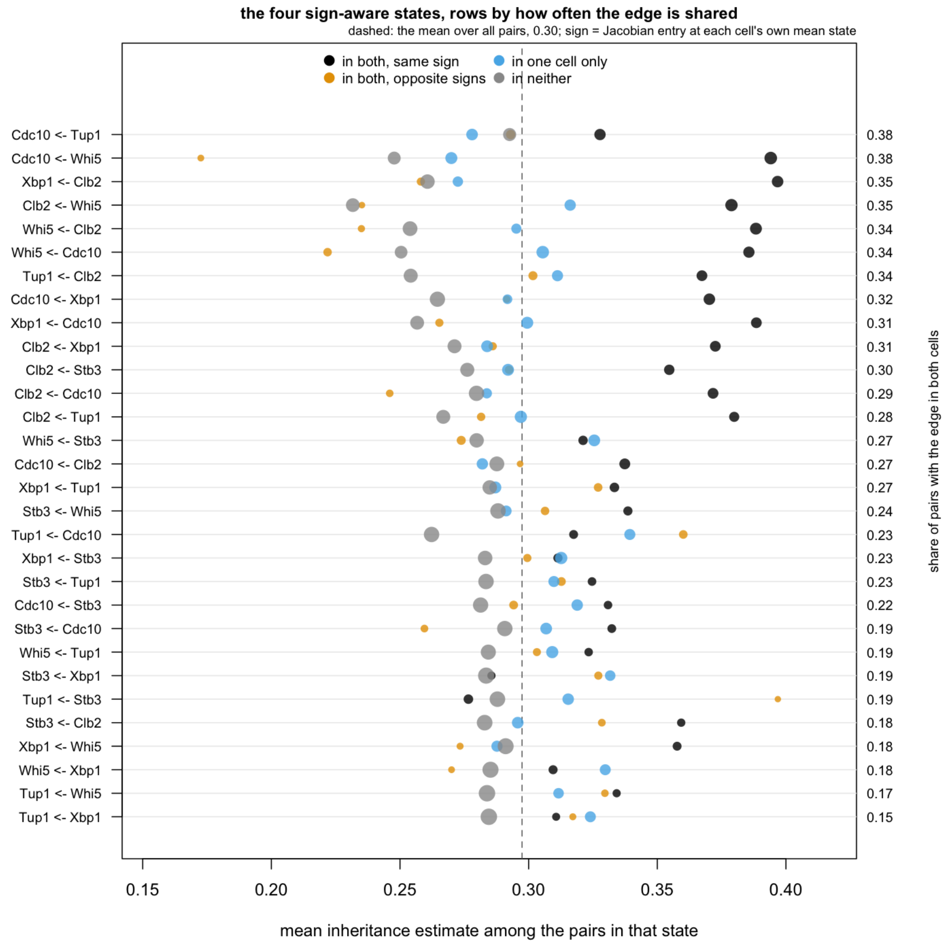


**Figure S8. Mean inheritance by the sign state of each edge.** One row per off-diagonal edge (written target ← regulator), ordered by the share of the 180 pairs in which both cells have the edge (right axis). Each point is the mean inheritance estimate over the pairs in one of four states, read from the sign of the edge's Jacobian entry at each cell's own mean state: in both cells with the same sign (black), in exactly one cell (blue), in neither (gray), or in both cells with opposite signs (orange); the point's size grows with the number of pairs in the state, and the dashed line is the mean over all pairs.

Figure S9 shows the individual pairs behind six rows of Figure S8, the six most frequently shared edges after Whi5 to Cdc10 (which is the right panel of Figure 8C), each drawn as in that panel with every pair's estimate jittered within its state, the state's mean as a bar, a Kruskal-Wallis $p$-value across the four states in the title, and a Wilcoxon $p$-value of the same-sign pairs against all others inside the panel. The ordering of the means is the same in every panel, with the same-sign pairs highest, but the evidence differs: for Clb2 to Xbp1, Whi5 to Clb2, Clb2 to Whi5, and Cdc10 to Whi5 the Kruskal-Wallis $p$ is at most $10^{-4}$ and the same-sign group is separated from the rest at $p \leq 10^{-4}$, for Clb2 to Tup1 the separation is weaker ($p = 0.003$), and for Tup1 to Cdc10, the single most frequently shared edge, the four states do not differ ($p = 0.5$), because its opposite-sign state is as common as its same-sign state and carries estimates just as high. That the most frequently shared edge is the one that does not track inheritance supports reading Tup1 to Cdc10 as a shared timing of two cycle-dependent signals rather than as a regulation that a daughter inherits.

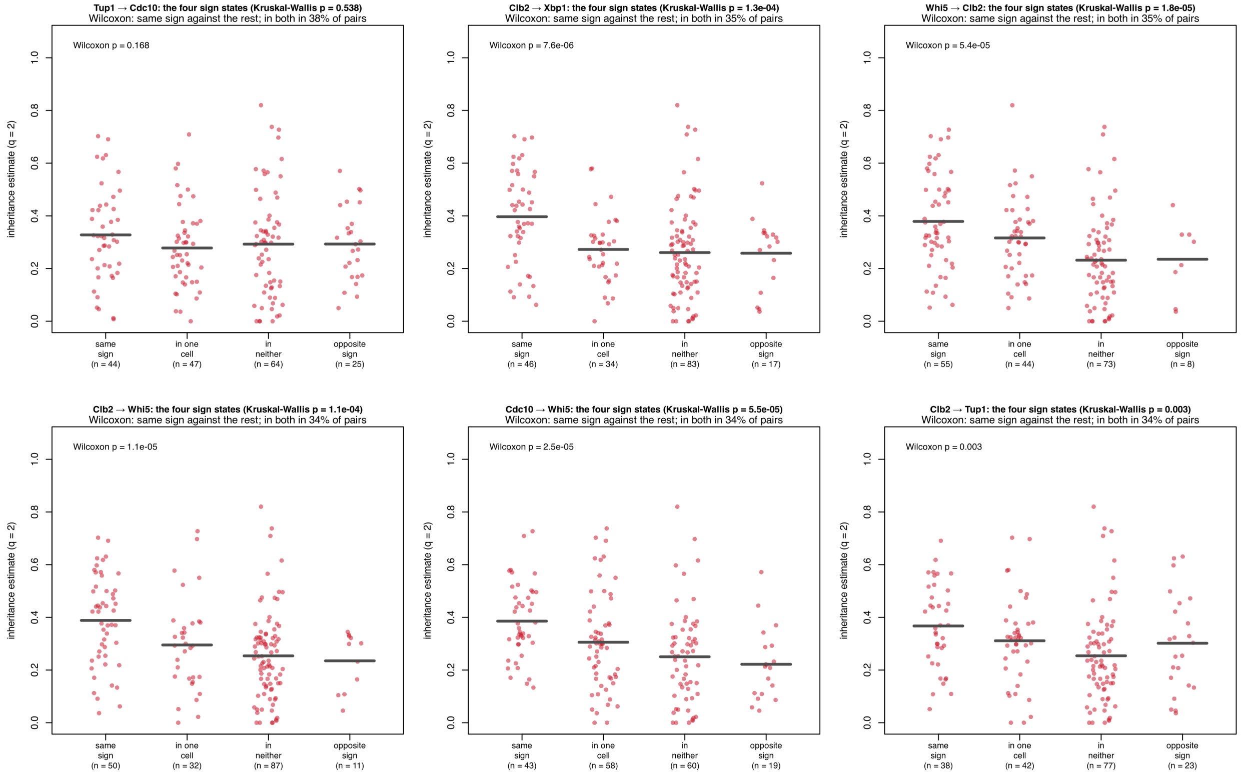


**Figure S9. The four sign states of mother-daughter pairs across the six most frequently shared edges after Whi5 → Cdc10.** Each panel is one edge, drawn as the right panel of Figure 8C: every pair's inheritance estimate is plotted within its "sign state," with the state's mean as a bar and the number of pairs below each state. The title gives the share of pairs with the edge in both cells and a Kruskal-Wallis $p$-value across the four states; the $p$-value inside each panel is a Wilcoxon rank-sum test of the same-sign pairs against all other pairs.

Figure S10 shows two edges from the bottom of Figure S8, Clb2 to Stb3 and Whi5 to Xbp1, each shared by 18% of pairs, which are of interest for the opposite reason: although few pairs share them, the pairs that do, with the same sign, have a mean inheritance of 0.36, as high as for the frequent edges, against 0.27 to 0.33 for the pairs in the other states. Because the same-sign groups hold only 18 and 22 pairs, neither edge reaches significance (Wilcoxon $p = 0.07$ and 0.10; Kruskal-Wallis $p = 0.2$ and 0.4), so these are candidates rather than findings. If they hold up in a larger lineage, they would be the more interesting kind of inherited edge, since an edge that most cells lack and that a daughter nevertheless shares with her mother is closer to a lineage-specific deviation than the consensus edges of Figure S9, and both link a cell-cycle protein to a stress repressor.

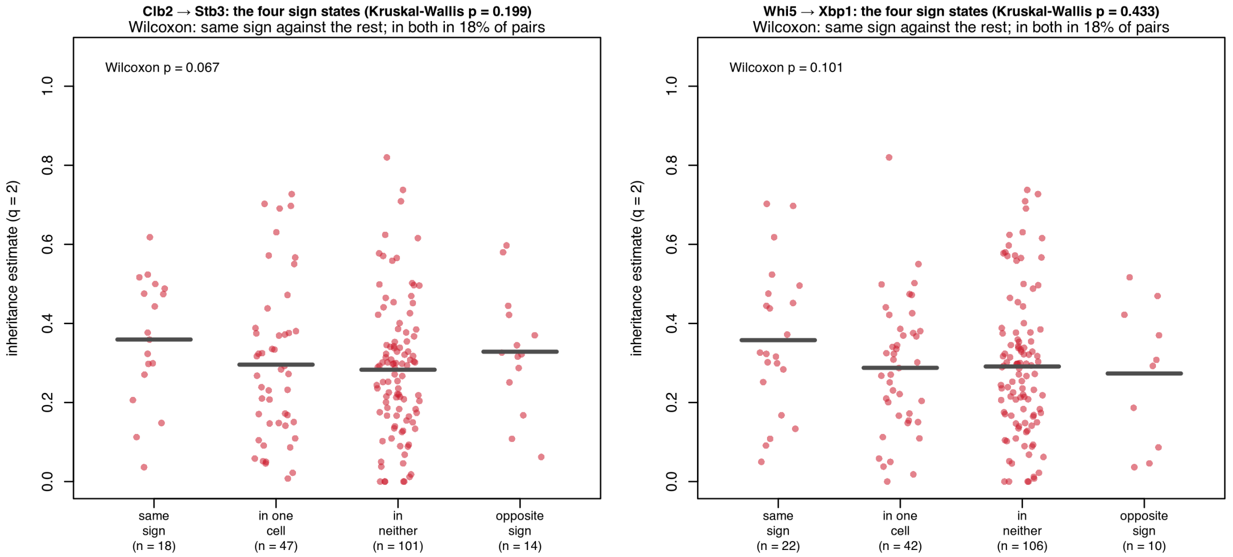


**Figure S10. The four "sign states" of two rarely shared edges, Clb2 → Stb3 and Whi5 → Xbp1.** Drawn as Figure S9. Each edge is present in both cells of 18% of the pairs; among those with the same sign, the mean inheritance is as high as for the most frequent edges, but the groups are small and neither edge reaches significance.

### H Proofs for theory in Appendix E

#### H.1 Proofs of Propositions 1 and 2, and the sharing rule at $q = 1$

Throughout, $j$ is fixed and dropped, $w_g = \sqrt{q}\mu_g$ with $\mu_S = 1$, and $X_{g,k}$, $\widetilde{\mathbf{y}}$ are as in Appendix B, so that (S3) reads $F(\beta) = \frac{1}{2}\|\widetilde{\mathbf{y}} - \sum_{g,k} X_{g,k}\beta_{g,k}\|_2^2 + \lambda \sum_{g,k} w_g \|X_{g,k}\beta_{g,k}\|_2$ with $\beta_{S,k} = \theta_k^{(S)}$ and $\beta_{c,k} = \delta_k^{(c)}$.

**Lemma 1 (Reduction).** *The loss in* (S3) *depends on* $(\theta^{(S)}, \delta^{(M)}, \delta^{(D)})$ *only through* $\theta_k^{(c)} = \theta_k^{(S)} + \delta_k^{(c)}$, *so* $(\widehat{\theta}^{(M)}, \widehat{\theta}^{(D)})$ *minimizes* $\sum_c \frac{1}{2}\|\widetilde{y}^{(c)} - \widetilde{\Psi}^{(c)}\theta^{(c)}\|_2^2 + \lambda\sqrt{q}\sum_k P_k(\theta_k^{(M)}, \theta_k^{(D)})$ *with the convex penalty*

$$P_k(u, v) = \min_{s \in \mathbb{R}^q} \|s\|_{S,k} + \mu_M\|u - s\|_{M,k} + \mu_D\|v - s\|_{D,k}, \tag{S10}$$

*and the scaffold and deviations at the solution are the minimizer* $s$ *in* (S10) *at* $(u, v) = (\widehat{\theta}_k^{(M)}, \widehat{\theta}_k^{(D)})$ *together with* $u - s$ *and* $v - s$.

*Proof.* The loss is a sum over cells of functions of $\theta_k^{(S)} + \delta_k^{(c)}$, and minimizing (S3) over $\theta^{(S)}$ first, for fixed $(\theta^{(M)}, \theta^{(D)})$, gives (S10), which is convex as the partial minimization of a jointly convex function. □

*Proof (of Proposition 1).* Let $X_{g,k} = Q_{g,k}R_{g,k}$ be a thin QR decomposition, which exists with $R_{g,k}$ invertible by the rank assumption, and set $b_{g,k} = R_{g,k}\beta_{g,k}$, so that $F(\beta) = \frac{1}{2}\|\widetilde{\mathbf{y}} - \sum_{g,k} Q_{g,k}b_{g,k}\|_2^2 + \lambda\sum_{g,k} w_g\|b_{g,k}\|_2$. This function is convex[4], so $b = 0$ is a minimizer if and only if 0 lies in its subdifferential at 0, that is, if and only if for every group there is a $z_{g,k}$ with $\|z_{g,k}\|_2 \le 1$ and $-Q_{g,k}^\top\widetilde{\mathbf{y}} + \lambda w_g z_{g,k} = 0$, which holds if and only if $\|Q_{g,k}^\top\widetilde{\mathbf{y}}\|_2 \le \lambda w_g$ for every group. Since $Q_{g,k}Q_{g,k}^\top$ is the projection onto the columns of $X_{g,k}$, $\|Q_{g,k}^\top\widetilde{\mathbf{y}}\|_2 = \|X_{g,k}(X_{g,k}^\top X_{g,k})^{-1}X_{g,k}^\top\widetilde{\mathbf{y}}\|_2 = \|(X_{g,k}^\top X_{g,k})^{-1/2}X_{g,k}^\top\widetilde{\mathbf{y}}\|_2$, and the condition is (S8). If the inequality is strict for every group and $b \neq 0$, then $F(b) - F(0) = -\widetilde{\mathbf{y}}^\top\sum_{g,k} Q_{g,k}b_{g,k} + \frac{1}{2}\|\sum_{g,k} Q_{g,k}b_{g,k}\|_2^2 + \lambda\sum_{g,k} w_g\|b_{g,k}\|_2 \ge \sum_{g,k}\left(\lambda w_g - \|Q_{g,k}^\top\widetilde{\mathbf{y}}\|_2\right)\|b_{g,k}\|_2 > 0$, so 0 is the unique minimizer. □

[4] These statements are derived from the Karush-Kuhn-Tucker condition of a convex problem without constraints [5], which for the group lasso reads: $b$ is a minimizer if and only if, for every group, $Q_{g,k}^\top(\widetilde{\mathbf{y}} - \sum_{g',k'} Q_{g',k'}b_{g',k'}) = \lambda w_g\, b_{g,k}/\|b_{g,k}\|_2$ when $b_{g,k} \neq 0$ and $\|Q_{g,k}^\top(\widetilde{\mathbf{y}} - \sum_{g',k'} Q_{g',k'}b_{g',k'})\|_2 \le \lambda w_g$ when $b_{g,k} = 0$ [31].

*Proof (of Proposition 2).* Fix $k$ and $(u,v)$, and let $s \in \mathbb{R}^q$. By the triangle inequality, $\mu_M \|u-s\|_{M,k} + \mu_D\|v-s\|_{D,k} \geq \mu_M\|u\|_{M,k} + \mu_D\|v\|_{D,k} - \big(\mu_M\|s\|_{M,k} + \mu_D\|s\|_{D,k}\big)$, and by the Cauchy-Schwarz inequality, $\mu_M\|s\|_{M,k} + \mu_D\|s\|_{D,k} \leq (\mu_M^2+\mu_D^2)^{1/2}\big(\|s\|_{M,k}^2 + \|s\|_{D,k}^2\big)^{1/2} = (\mu_M^2+\mu_D^2)^{1/2}\|s\|_{S,k} \leq \|s\|_{S,k}$. Hence the objective in (S10) at $s$ is at least $\mu_M\|u\|_{M,k} + \mu_D\|v\|_{D,k}$, its value at $s=0$, and because $\mu_M^2+\mu_D^2 < 1$ the last inequality is strict for $s \neq 0$, so $s=0$ is the unique minimizer and $P_k(u,v) = \mu_M\|u\|_{M,k} + \mu_D\|v\|_{D,k}$ (at $\mu_M^2+\mu_D^2 = 1$, $s=0$ is still a minimizer, so two separate fits still minimize (S3)). By Lemma 1, $(\widehat{\theta}^{(M)}, \widehat{\theta}^{(D)})$ then minimizes $\sum_c \big[\frac{1}{2}\|\widetilde{y}^{(c)} - \widetilde{\Psi}^{(c)}\theta^{(c)}\|_2^2 + \lambda\sqrt{q}\mu_c \sum_k \|\theta_k^{(c)}\|_{c,k}\big]$, which separates into the two single-cell problems of the statement. □

*Proof (of Proposition 3).* At $q=1$, $\|v\|_{c,k} = a_{c,k}|v|$ and $\|s\|_{S,k} = a_{S,k}|s|$, so (S10) is the weighted absolute-deviation problem $\min_s W_S|s-0| + W_M|s-u| + W_D|s-v|$ with weights $W_S = a_{S,k}$, $W_M = \mu_M a_{M,k}$, and $W_D = \mu_D a_{D,k}$ at the points 0, $u$, and $v$, whose minimizers are the weighted medians of these three points. Take $0 < u \leq v$; the other same-sign case is symmetric. The cumulative weights in the order $0, u, v$ are $W_S$, $W_S + W_M$, and $W = W_S + W_M + W_D$, so $s=0$ is a minimizer if and only if $W_S \geq W/2$, that is $a_{S,k} \geq \mu_M a_{M,k} + \mu_D a_{D,k}$, and the unique minimizer when the inequality is strict. When $W_S < W/2$, the minimizer is $u$ if $W_S + W_M \geq W/2$, that is $W_D \leq W_S + W_M$, and $v$ otherwise, and in either case it is one of the two coefficients, whose cell then has a zero deviation. At $\mu_M = \mu_D = 1$, $W_c = a_{c,k} \leq a_{S,k} = W_S$, so $W_D \leq W_S + W_M$ holds and the scaffold is $u$, the coefficient of smaller magnitude; for $u < 0 < v$ the cumulative weights in the order $u, 0, v$ are $W_M$ and $W_M + W_S$, and $W_M \leq W_S + W_D$ together with $W_D \leq W_S + W_M$ places the median at 0. Finally, $a_{S,k}^2 = a_{M,k}^2 + a_{D,k}^2$ gives $a_{S,k} < a_{M,k} + a_{D,k}$ when both are positive and $a_{S,k} \geq (a_{M,k} + a_{D,k})/\sqrt{2}$ by $(a_{M,k} - a_{D,k})^2 \geq 0$, which are the bounds quoted before the proof. □

#### H.2 Proof of Proposition 4

*Proof (of Proposition 4).* Write $\Phi^{(c)} = \exp(hA^{(c)})$, so that (6) reads $\tau_h = 1 - \|\Phi^{(M)} - \Phi^{(D)}\|_F / (\|\Phi^{(M)} - I\|_F + \|\Phi^{(D)} - I\|_F)$.

(a) From the series of the matrix exponential, $\Phi^{(c)} - I = hA^{(c)} + E_h^{(c)}$ with $\|E_h^{(c)}\|_F \leq \sum_{m\geq 2} h^m \|A^{(c)}\|_F^m/m! \leq h^2\|A^{(c)}\|_F^2 e^{h\|A^{(c)}\|_F}$, so $\|E_h^{(c)}\|_F = O(h^2)$. By the triangle inequality, the numerator of $1-\tau_h$ is $h\|A^{(M)} - A^{(D)}\|_F + O(h^2)$ and the denominator is $h(\|A^{(M)}\|_F + \|A^{(D)}\|_F) + O(h^2)$. Because $\|A^{(M)}\|_F + \|A^{(D)}\|_F > 0$, the ratio equals $\|A^{(M)} - A^{(D)}\|_F / (\|A^{(M)}\|_F + \|A^{(D)}\|_F) + O(h)$, which is the claim.

(b) Expanding the squared norm, $\|A^{(M)} - A^{(D)}\|_F^2 = \|A^{(M)}\|_F^2 + \|A^{(D)}\|_F^2 - 2\,\mathrm{tr}(A^{(M)\top}A^{(D)})$. With $\|A^{(M)}\|_F = \|A^{(D)}\|_F = a$ and a zero trace term, the numerator of $1-\tau_0$ is $\sqrt{2}a$ and the denominator is $2a$, so $\tau_0 = 1 - \sqrt{2}/2 = 1 - 1/\sqrt{2}$.

□

#### References


1. Amon, A., Tyers, M., Futcher, B., Nasmyth, K.: Mechanisms that help the yeast cell cycle clock tick: G2 cyclins transcriptionally activate G2 cyclins and repress G1 cyclins. Cell **74**(6), 993–1007 (1993)
2. Argüello-Miranda, O., Marchand, A.J., Kennedy, T., Russo, M.A., Noh, J.: Cell cycle–independent integration of stress signals by Xbp1 promotes non-G1/G0 quiescence entry. Journal of Cell Biology **221**(1), e202103171 (2021)
3. Bergen, V., Lange, M., Peidli, S., Wolf, F.A., Theis, F.J.: Generalizing RNA velocity to transient cell states through dynamical modeling. Nature Biotechnology **38**(12), 1408–1414 (2020)
4. Biddy, B.A., Kong, W., Kamimoto, K., Guo, C., Waye, S.E., Sun, T., Morris, S.A.: Single-cell mapping of lineage and identity in direct reprogramming. Nature **564**(7735), 219–224 (2018)
5. Boyd, S., Vandenberghe, L.: Convex Optimization. Cambridge University Press, Cambridge (2004)
6. Brunton, S.L., Proctor, J.L., Kutz, J.N.: Discovering governing equations from data by sparse identification of nonlinear dynamical systems. Proceedings of the National Academy of Sciences **113**(15), 3932–3937 (2016)
7. Chen, K.C., Calzone, L., Csikasz-Nagy, A., Cross, F.R., Novak, B., Tyson, J.J.: Integrative analysis of cell cycle control in budding yeast. Molecular Biology of the Cell **15**(8), 3841–3862 (2004)
8. Chen, S., Shojaie, A., Witten, D.M.: Network reconstruction from high-dimensional ordinary differential equations. Journal of the American Statistical Association **112**(520), 1697–1707 (2017). `https://doi.org/10.1080/01621459.2016.1229197`, `https://doi.org/10.1080/01621459.2016.1229197`, pMID: 29618851
9. Huguet, G., Magruder, D.S., Tong, A., Fasina, O., Kuchroo, M., Wolf, G., Krishnaswamy, S.: Manifold interpolating optimal-transport flows for trajectory inference. In: Advances in Neural Information Processing Systems. vol. 35 (2022)
10. Jindal, K., Adil, M.T., Yamaguchi, N., Yang, X., Wang, H.C., Kamimoto, K., Rivera-Gonzalez, G.C., Morris, S.A.: Single-cell lineage capture across genomic modalities with CellTag-multi reveals fate-specific gene regulatory changes. Nature Biotechnology **42**, 946–959 (2024)
11. Kennedy, T., Yalcinkaya, B., Ramakanth, S., Neupane, S., Tadić, N., Buchler, N.E., Argüello-Miranda, O.: Deep learning–driven imaging of cell division and cell growth across an entire eukaryotic life cycle. Molecular Biology of the Cell **36**(6), ar74 (2025)
12. La Manno, G., Soldatov, R., Zeisel, A., Braun, E., Hochgerner, H., Petukhov, V., Lidschreiber, K., Kastriti, M.E., Lönnerberg, P., Furlan, A., et al.: RNA velocity of single cells. Nature **560**(7719), 494–498 (2018)
13. Li, J., Pan, X., Yuan, Y., Shen, H.B.: TFvelo: gene regulation inspired RNA velocity estimation. Nature Communications **15**, 1387 (2024)
14. Liko, D., Conway, M.K., Grunwald, D.S., Heideman, W.: Stb3 plays a role in the glucose-induced transition from quiescence to growth in Saccharomyces cerevisiae. Genetics **185**(3), 797–810 (2010)
15. Mai, B., Breeden, L.: Xbp1, a stress-induced transcriptional repressor of the Saccharomyces cerevisiae Swi4/Mbp1 family. Molecular and Cellular Biology **17**(11), 6491–6501 (1997)
16. McMurray, M.A., Bertin, A., Garcia, G., Lam, L., Nogales, E., Thorner, J.: Septin filament formation is essential in budding yeast. Developmental cell **20**(4), 540–549 (2011)
17. Meier, L., van de Geer, S., Bühlmann, P.: The group lasso for logistic regression. Journal of the Royal Statistical Society: Series B (Statistical Methodology) **70**(1), 53–71 (2008)
18. Messenger, D.A., Bortz, D.M.: Weak SINDy: Galerkin-based data-driven model selection. Multiscale Modeling & Simulation **19**(3), 1474–1497 (2021)
19. Örd, M., Venta, R., Möll, K., Valk, E., Loog, M.: Cyclin-specific docking mechanisms reveal the complexity of M-CDK function in the cell cycle. Molecular Cell **75**(1), 76–89 (2019)
20. Qiu, W., Neo, S.P., Yu, X., Cai, M.: A novel septin-associated protein, Syp1p, is required for normal cell cycle-dependent septin cytoskeleton dynamics in yeast. Genetics **180**(3), 1445–1457 (2008)
21. Reinbold, P.A.K., Gurevich, D.R., Grigoriev, R.O.: Using noisy or incomplete data to discover models of spatiotemporal dynamics. Physical Review E **101**(1), 010203 (2020)
22. Schwarz, G.: Estimating the dimension of a model. The Annals of Statistics **6**(2), 461–464 (1978)
23. Setty, M., Kiseliovas, V., Levine, J., Gayoso, A., Mazutis, L., Pe'er, D.: Characterization of cell fate probabilities in single-cell data with Palantir. Nature Biotechnology **37**(4), 451–460 (2019)

24. Simon, N., Tibshirani, R.: Standardization and the group lasso penalty. Statistica Sinica **22**(3), 983–1001 (2012), `http://www.jstor.org/stable/24309971`
25. Skotheim, J.M., Di Talia, S., Siggia, E.D., Cross, F.R.: Positive feedback of G1 cyclins ensures coherent cell cycle entry. Nature **454**(7202), 291–296 (2008)
26. Smith, R.L., Johnson, A.D.: Turning genes off by Ssn6-Tup1: A conserved system of transcriptional repression in eukaryotes. Trends in Biochemical Sciences **25**(7), 325–330 (2000)
27. Street, K., Risso, D., Fletcher, R.B., Das, D., Ngai, J., Yosef, N., Purdom, E., Dudoit, S.: Slingshot: cell lineage and pseudotime inference for single-cell transcriptomics. BMC Genomics **19**, 477 (2018)
28. Tibshirani, R., Bien, J., Friedman, J., Hastie, T., Simon, N., Taylor, J., Tibshirani, R.J.: Strong rules for discarding predictors in lasso-type problems. Journal of the Royal Statistical Society Series B: Statistical Methodology **74**(2), 245–266 (2012)
29. Tronstad, M., Karlsson, J., Dahlin, J.S.: MultistageOT: Multistage optimal transport infers trajectories from a snapshot of single-cell data. Proceedings of the National Academy of Sciences **122**(50), e2516046122 (2025)
30. Weinreb, C., Rodriguez-Fraticelli, A., Camargo, F.D., Klein, A.M.: Lineage tracing on transcriptional landscapes links state to fate during differentiation. Science **367**(6479), eaaw3381 (2020)
31. Yuan, M., Lin, Y.: Model selection and estimation in regression with grouped variables. Journal of the Royal Statistical Society: Series B (Statistical Methodology) **68**(1), 49–67 (2006)
